# Gene-transcription factor regulatory networks implicate primary cilia in the evolution of vertebrate sex determination and expand models of epigenetic regulation

**DOI:** 10.1101/2025.11.21.689810

**Authors:** Thea B. Gessler, Dean C. Adams, Nicole Valenzuela

**Affiliations:** Department of Ecology, Evolution, and Organismal Biology, Iowa State University, Ames, Iowa, USA; Genetics and Genomics Program, Iowa State University, Ames, Iowa, USA

## Abstract

The genetic architecture underlying diverse vertebrate sex-determining systems remains elusive despite evidence of changes in upstream regulators and downstream mediators. Here we modeled species-specific regulatory networks of gonadal development for turtles with contrasting mechanisms [*Apalone spinifera –* ZZ/ZW genotypic sex determination (GSD), and *Chrysemys picta* – temperature-dependent sex determination (TSD)] using matched time-course sampling. We uncovered key steps in the evolutionary transition in sex determination by testing for conservation or divergence of network modular components. Specifically, we tested these alternative hypotheses: first, transcription factor (TF) hubs and their target genes are conserved between species (null H0); second, the same TF hub acquired a new set of target genes in a species, retaining or not ancestral functions (H1 and variants); third, a new TF hub takes over the regulation of the former gene targets of an ancestral TF (H2); and finally, complete overhaul occurs where both ancestral TF hubs and their target genes were replaced in a species (H3). Results implicate primary cilia as integrators of environmental signals underlying TSD, as known thermosensitive TSD components (e.g., calcium-redox, pSTAT3, *Wnt*/*Rspo1*/*B-catenin*, *Dhh*) are linked to primary cilia. TFs that evolved between species also regulate primary cilia and point to key changes in their sensory machinery that accompanied TSD-GSD transitions (e.g., calcium/ion channels or membrane transport components in *Chrysemys* versus structural elements and ciliogenesis in *Apalone*). This novel Primary Cilia Integration hypothesis expands current models of epigenetic regulation of turtle sexual development, the evolution of plasticity versus canalization, and warrants functional validation.

## Introduction

Vertebrate sex determination provides a compelling example of a developmental mechanism that diverges greatly despite sharing highly conserved gene players (Morrish and Sinclair 2002; Bachtrog, et al. 2014; Capel 2017; Stöck, et al. 2021), a case of developmental systems drift (True and Haag 2001). Specifically, vertebrate sex-determining mechanisms (SDMs) span a spectrum from strict environmental sex determination (ESD) to strongly canalized genotypic sex determination (GSD) (Valenzuela 2008c; Tree of Sex Consortium, et al. 2014). SDMs are governed by genes recycled frequently among taxa, but their position or regulation in the network is shuffled repeatedly by evolution, such as occurred for *Dmrt1* (Smith, et al. 2009; Cui, et al. 2017; Ge, et al. 2017), *Sox9* (Morais da Silva, et al. 1996), and *Aromatase* (Navarro-Martin, et al. 2011), among many others. Importantly, our full understanding of the rewiring of the genetic network underlying the diversity of vertebrate SDMs is obscured in part because much of our knowledge still relies on mouse and human models (Bachtrog, et al. 2014).

Turtles are a vertebrate group particularly suited to unravel developmental network divergence as they showcase temperature-dependent sex determination (TSD)—a type of ESD, plus XX/XY and ZZ/ZW sex chromosomal systems of GSD, that evolved multiple times independently accompanied by accelerated molecular evolution of sexual development genes (Valenzuela and Adams 2011a; Consortium 2014; Literman, Burrett, et al. 2018; Bista and Valenzuela 2020). These natural experiments enable comparative analyses between closely related lineages with and without plastic sexual development to uncover the shifts in genetic architecture underlying these transitions (Valenzuela 2008a; Literman, Burrett, et al. 2018) aided by growing ‘omic resources in turtles (Badenhorst, et al. 2015; Czerwinski, et al. 2016; Gessler, et al. 2023a). Active research is devoted to identifying the elusive environmental sensor(s) that distinguishes TSD from GSD. Recent potential candidates include calcium and redox (CaRe) status sensors (Castelli, et al. 2020a), and epigenetic histone modifications driving the temperature-specific activation or repression of *Dmrt1* (Ge, et al. 2017; Weber, et al. 2020a), a masculinizing gene implicated in a GSD turtle as well (Sun, et al. 2017). However, broader changes have accrued between TSD and GSD networks beyond these few candidate elements. Indeed, comparisons across turtles and other vertebrates revealed that transcriptional heterochrony contributes to the divergence in SDMs (Valenzuela 2008c; Valenzuela, et al. 2013; Capel 2017; Mizoguchi and Valenzuela 2020), such that agnostic genome-wide investigations should substantially illuminate these evolutionary events.

Divergence of developmental programs underlying phenotypic evolution often involve the evolution of gene regulatory network (GRN) components, such as cis-regulatory elements (CREs) that change the interaction between transcription factors (TFs) and the genes they regulate (Carroll 2008). This is true for GRNs controlling animal sexual development, which at a broad taxonomic scale appear quite evolutionarily labile at the upstream trigger and more conserved in the downstream TFs (Williams and Carroll 2009). TFs may regulate a multitude of genes in a GRN, acting as TF hubs, and have pleiotropic effects in multiple biological processes. Thus, because changes in hubs can lead to profound phenotypic changes with potentially detrimental fitness effects (Carroll 2008), highly connected and central hubs are expected to evolve at slower rate than peripheral elements of the GRN (Helsen, et al. 2019) [although adaptive evolution via changes to central hubs may be possible (Koubkova-Yu, et al. 2018; Helsen, et al. 2019)], whereas changes can accumulate more gradually via shifts in gene target identity (Carroll 2008).

Here we constructed and compared gene-transcription factor interaction networks for two turtle species with contrasting SDMs: the painted turtle *Chrysemys picta*, a TSD representative used as proxy for the ancestral condition in turtles (Valenzuela and Adams 2011a; Sabath, et al. 2016; Bista and Valenzuela 2020), and the spiny softshell turtle *Apalone spinifera*, a GSD species with an evolutionarily derived ZZ/ZW mechanism (Badenhorst, et al. 2013; Sabath, et al. 2016; Bista and Valenzuela 2020), acknowledging that evolution has occurred in both lineages since their split ∼175 Mya (timetree.org). This approach allowed us to build testable models about the molecular circuitry of TSD and GSD, how it evolved in these distinct lineages, and to identify candidate elements for a putative role as key TSD or GSD regulators, including the involvement of primary cilia [which are organelles that nearly all cell types use to sense the context of cues from the external environment or internal signaling pathways (Mill, et al. 2023)] in the evolution of vertebrate sex determination. Species will be referred to by their genus name hereafter. Specifically, we tested the following non-mutually exclusive hypotheses about evolutionary patterns of turtle sexual development networks, by examining changes between species in TF hubs (i.e., TFs regulating many genes in the network), and their target genes (Fig 1). Of note, we use the terms hub and TF interchangeably, as the TFs examined here act as hubs within the subnetworks of the genes they target (the focus of much of our comparisons), while recognizing that in the context of broader networks some TFs may be more interconnected (i.e., hub-like) than others:

**Fig 1.**
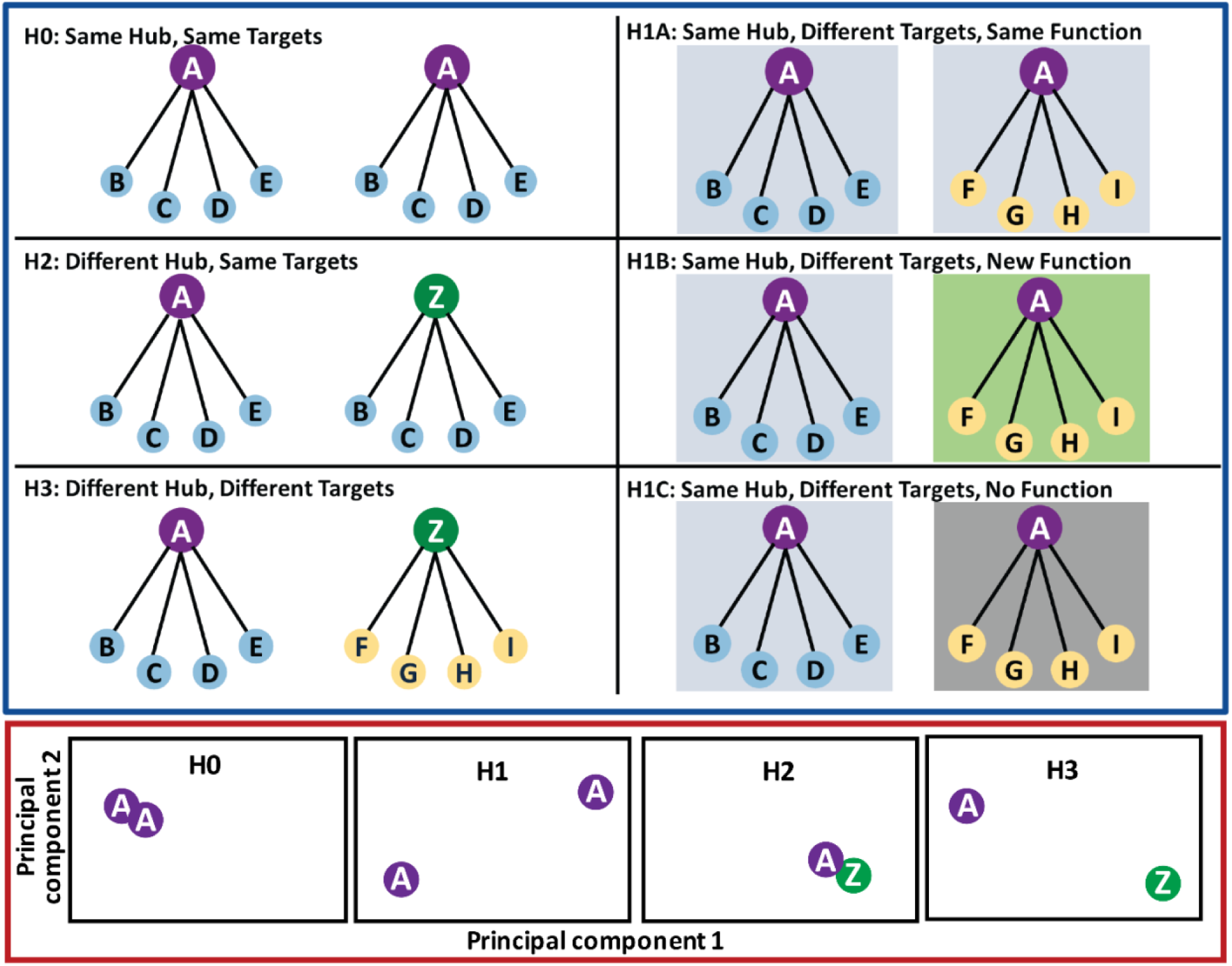
Illustration of predictions from the hypotheses of the evolution of vertebrate sex determination by shifts in the interactions of transcription factors and their predicted target genes. Top three rows encircled in blue: (H0) null hypothesis, i.e. network/module conservation, (H1) upstream conservation, downstream divergence, (H2) upstream divergence, downstream conservation, and (H3) full network/module overhaul. Bottom row encircled in red: depiction of the pattern of TF clustering in regulome space (principal components plot capturing differences in targeting patterns of TF hubs) expected under hypotheses H0, H1, H2, and H3 (see text for details).

### H0: Same hub, same targets hypothesis

Neither the hub TFs nor their target genes vary between species. If this null hypothesis is supported, it would indicate evolutionary conservation of network wiring between TSD and GSD. Partial support of this hypothesis for only some TFs would expose network connections under strong stabilizing selection. In contrast, such a finding for the entire network would be surprising given that these taxa split 175 Mya (timetree.org) and *Apalone*’s softshell turtle family shifted from TSD to GSD.

### H1A: Same hub directs same function with new targets

A conserved TF hub recruits a novel set of target genes that take over the regulation of the ancestral function. Support for H1A would uncover TFs subject to developmental systems drift (True and Haag 2001) because the TF continues to direct the same function (determined by finding a conserved function of the targets in an overrepresentation test), but the gene targets implementing that function have changed.

### H1B: Same hub directs different function with new targets

A conserved TF hub recruits a novel set of target genes, but these targets regulate a new function. Such a change could occur *via* natural selection for a new function via the rewiring of the TF’s target genes (in which case an overrepresentation test would reveal species-specific functions for each set of gene targets).

### H1C: Same hub, new targets with no function

The TF hub remains conserved, but it regulates new target genes that have no detectable function as determined via overrepresentation tests (i.e., pseudofunctionalization). This pattern could result from genetic drift, perhaps when the original function regulated by a TF is released from selection by changes elsewhere in the network, facilitating gains or losses of TF binding sites (TFBSs) irrespective of their functional potential.

### H2: New hub, same targets

A new TF hub takes over the regulation of ancestral gene targets. If supported, H2 would uncover the evolution of upstream regulation of network modules underlying sexual development that could occur by natural selection or developmental systems drift. This would be demonstrated by finding a similar set of regulatory targets and functions for non-orthologous TFs between species.

### H3: New hub, different targets

H3 implies the overhaul of the sexual development network either in its entirety, or for some of its modules, and would be supported if major differences are found in the identity of TF hubs, their connectivity, and their target genes, which could be carrying out either the same function as the ancestral hub and targets, a new function (neofunctionalization), or no function (pseudofunctionalization).

## Results

### Putative core network components of turtle sexual development

Models of gene-TF interaction networks were constructed using PANDA (Glass, et al. 2013; Schlauch, et al. 2017) with matched-stage RNA-seq data from a previous study of *Chrysemys* and *Apalone* embryos (Gessler, et al. 2023a) (trunks at stage 9, adrenal kidney gonad complexes at stages 12 and 15, and gonads at stages 19 and 22), transcription binding sites from their reference genomes(Badenhorst, et al. 2015; Bista, et al. 2024), and vertebrate protein-protein interaction data (Szklarczyk, et al. 2019), which yielded two networks for *Chrysemys* (Cpi-Female-31°C and Cpi-Male-26°C), and four networks for *Apalone* (Asp-Female-26°C, Asp-Female-31°C, Asp-Male-26°C, and Asp-Male-31°C). We will refer to these networks as Cpi-FPT, Cpi-MPT, Asp-F26, Asp-M26, Asp-F31, and Asp-M31 hereafter. For *Chrysemys* and *Apalone*, all network pairs within species were nearly identical when analyzed for differential edges (an edge is a network vertex connecting a gene to a TF, representing the presence of an interaction), differential targeting patterns (i.e. gene in-degree and TF out-degree patterns, which describe how many edges point to a gene, and how many edges point from a TF to various genes, respectively), and overall network differences at the full matrix level (i.e., *Chrysemys*: Cpi*-*MPT vs Cpi-FPT; *Apalone*: Asp-M26 vs Asp-F26, Asp-M31 vs Asp-F31, Asp-M26 vs Asp-M31, and Asp-F26 vs Asp-F31). Namely, no differential edges or differences in targeting patterns were detected after Benjamini-Hochberg correction, and the sum of squares calculation assessing overall network differences was also non-significant (p > 0.2 in all cases, Table 1). The same was true if networks were built based on only differentially expressed genes. This overall similarity among networks at the global level is likely due to the small size of the dataset and the pooling of gene expression data from five developmental stages. Thus, this coarse approach identified conserved developmental signals common to all conditions and embryonic stages (both within and between species) that represent presumptive core components of turtle developmental processes to be functionally validated in future studies. Instead, the subtle sex- or temperature-specific differences that were too weak to be detected with this method (masked by the broad overall similarities), were revealed by quantitative hypothesis testing. This included 89 out of 148 turtle TFs from the full networks that were expressed in the time-course transcriptomes of both *Chrysemys* and *Apalone* and thus permitted further evolutionary analyses between species.

**Table 1.** Sum of squares result from the overall network comparison. Top three rows encircled in blue: (H0) null hypothesis, i.e. network/module conservation, (H1) upstream conservation, downstream divergence, (H2) upstream divergence, downstream conservation, and (H3) full network/module overhaul. Bottom row encircled in red: depiction of the pattern of TF clustering in regulome space (principal components plot capturing differences in targeting patterns of TF hubs) expected under hypotheses H0, H1, H2, and H3 (see text for details).

| Network Comparison | Sum of Squares | Pvalue |
| --- | --- | --- |
| *Chrysemys* 26°C vs 31°C | 330214.43 | 0.677 |
| *Apalone* Female 26°C vs Male 26°C | 226941.63 | 0.376 |
| *Apalone* Female 31°C vs Male 31°C | 264538.08 | 0.224 |
| *Apalone* Female 26°C vs Female 31°C | 146894.62 | 0.853 |
| *Apalone* Male 26°C vs Male 31°C | 237478.87 | 0.329 |

### Principal components and trajectory analysis

We employed principal components (PC) analysis on the edge weights representing the genes targeted by each TF to more clearly discern the patterns present in these hyperdimensional ‘omics data, after retaining only genes and TFs shared between species across all six networks, transforming negative edge-weights to zero to denote a lack of relationship, and excluding negligibly expressed TFs with average expression <1.5 log_2_(TPM) [TPM = transcripts per million base pairs]. Next, the principal components were subjected to a modified multivariate trajectory analysis (Adams and Collyer 2009) to test for differences in the regulatory targeting patterns of orthologous TFs, by comparing the length of the vector that connects centroids of orthologous TFs in regulome space (which measures how similar or different TFs are relative to one another in the genes they are predicted to regulate between species) to the length predicted under the null hypothesis (H0) that there were no differences between species (Fig. 2A-C). PC1 and PC2 captured 25.2% and 5.1% of the variation, respectively (Fig 2D), while 888 PCs explained all the variance, reflecting the high complexity of these networks and the many factors that contribute to their variation.

**Fig 2.**
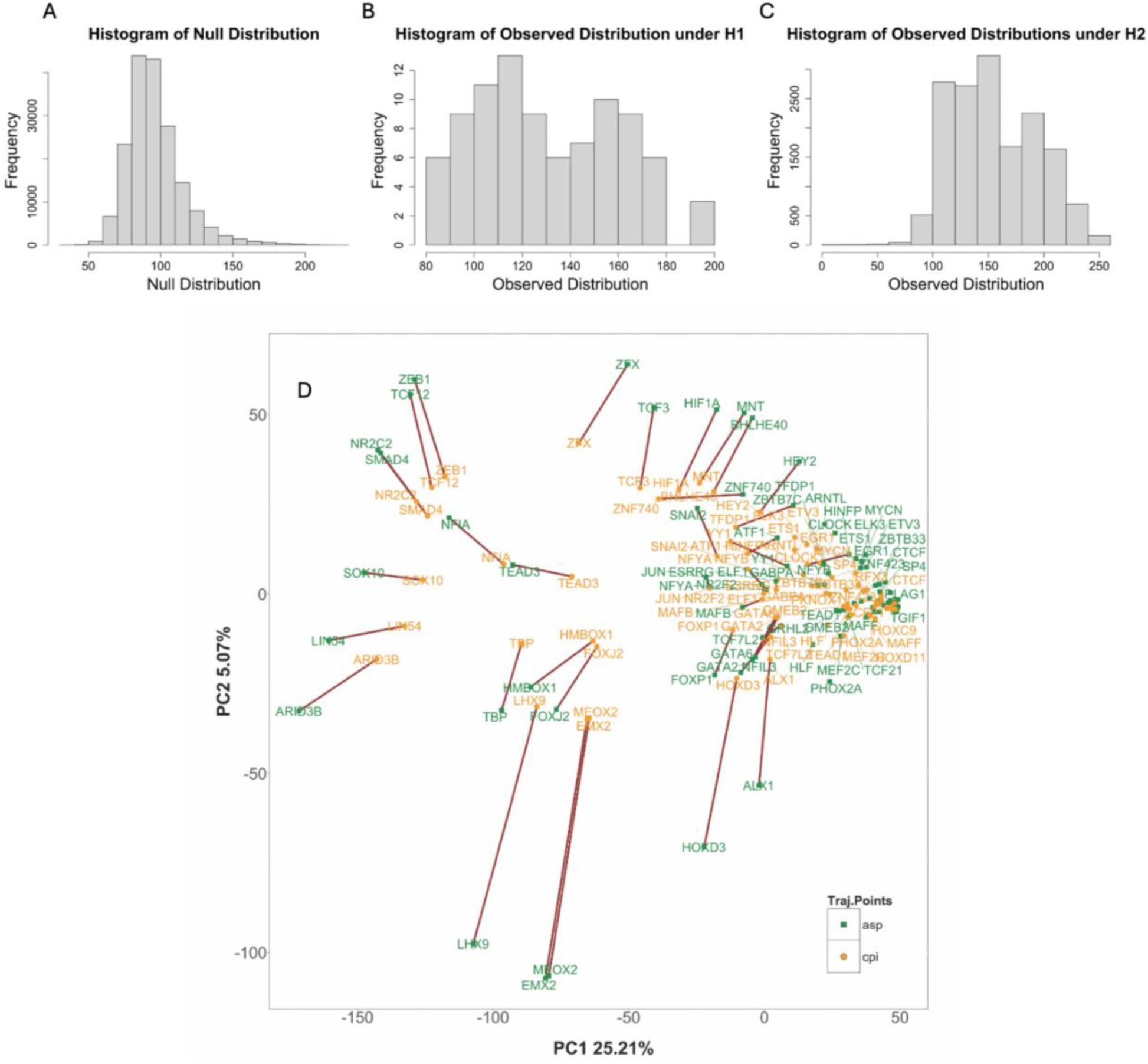
Trajectory analysis of the evolution of 89 transcription factors between *Chrysemys* and *Apalone*. Null (A) and observed (B and C) empirical distributions of distances (trajectory magnitudes) between orthologous TFs (B) and non-orthologous TFs (C) in regulome space (D). Trajectory magnitudes are based on the edge weights between TFs and their gene targets which are represented as effect sizes (Z scores). The distribution in panel B was used to test hypothesis H1, i.e. whether trajectories between orthologous TFs were longer than expected under the null hypothesis H0 (that no evolution occurred between these two turtle lineages over 175 my). The distribution in panel C was used to test hypothesis H2, i.e., whether trajectories between non-orthologous TFs were shorter than expected under the null hypothesis H0, which would indicate that non-orthologous TFs converged between species on a common set of gene targets. (D) Greater divergence between species in the gene targeting patterns of orthologous TFs is denoted by longer vectors (greater Euclidean distance in the principal components regulome space) between *Apalone* and *Chrysemys*. Thicker brown lines indicate significantly longer trajectories than expected, whereas thinner gray lines indicate trajectories of length expected by chance (supporting H0).

Results from the multivariate trajectory analysis on this principal components’ space (the regulome space) revealed 50 of 89 TFs that supported the null hypothesis H0 (same hub, same targets) as they were conserved in their regulatory targets (Fig 2A). Hypothesis H1 (same hub, different targets) (Fig 2B) was supported by the remaining 39 TFs that significantly diverged from their ortholog in the identity of genes targeted or in the strength of their connection (Fig 2D). Next, we functionally annotated these 39 TFs and found that 5 TFs (ARID3B, EMX2, LHX9, LIN54, and MEOX2) exhibited significant overrepresentation test results across all 6 networks (FDR of 0.05), enabling full cross-species comparison of their semantic similarity for hypotheses testing of the three alternative explanations of their divergence in regulome space (Table 2). Semantic similarity considers the gene ontology terms and graph structure to assess differences in functional annotation(Zhao and Wang 2018). We analyzed these 5 TFs to determine whether the functional role of their new targets remained conserved despite the turnover in target identity (hypothesis: H1A), whether a new functional role was acquired (hypothesis H1B), or whether a clear function was lost (hypothesis H1C) (Fig 1), and found ARID3B supports H1A, while EMX2, LHX9, LIN54, and MEOX2 support H1B. Support for hypothesis H2 that a non-orthologous TF took over the regulation of an ancestral TF hub, was found for a single TF in the GSD *Apalone* (ZBED1) that is closer in regulome space than expected under the null H0 to 12 TFs in the TSD *Chrysemys* (BACH2, BCL6, CTCF, GLI2, GLIS3, IRF1, MTF1, NR2F6, PKNOX2, RXRA, TGIF1, ZNF410). Intriguingly, ZBED1 orthologs were not significantly farther apart than expected (failed to reject H0), thus, ortholog targeting patterns did not differ between species. Combined, these results suggests that *Apalone* ZBED1 has acquired a new regulatory role without giving up its ancestral regulatory role.

**Table 2.**
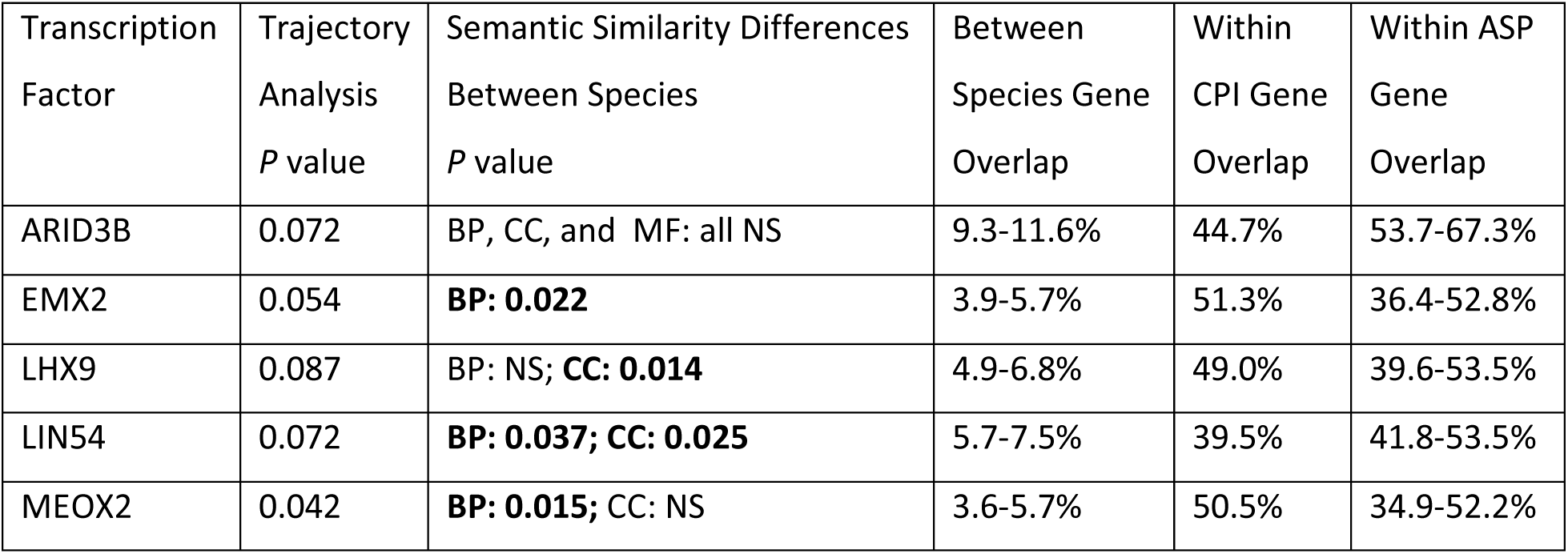
Semantic similarity in biological process (BP), cellular component (CC) and molecular function (MF) for transcription factors that evolved significant differences in targeting patterns between *Chrysemys* and *Apalone* as determined by trajectory analysis of the principal components regulome space. Semantic similarity indicates how similar the sets of gene ontology terms annotated to a TF were when compared between species while accounting for hierarchical graph structure. P-values are provided for TFs with significant differences in their semantic similarity between species at α = 0.1 which corresponded to the top 5% of TFs who occupied the upper mode of the observed distribution (Fig 2B). The degree of overlap in the sets of genes regulated by each TF is also provided, including: the range of gene overlap for all *Chrysemys* vs all *Apalone* (Between Species), *Chrysemys* MPT vs *Chrysemys* FPT (Within CPI), and the range of gene overlap for all *Apalone* vs all *Apalone* (Within ASP) to illustrate the degree of change in gene targeting patterns between species. See the Supplementary Materials for the full list of overrepresentation results. NS = Non-significant.

| Transcription Factor | Trajectory Analysis<br>*P* value | Semantic Similarity Differences<br>Between Species<br>*P* value | Between Species Gene Overlap | Within CPI Gene Overlap | Within ASP Gene Overlap |
| --- | --- | --- | --- | --- | --- |
| ARID3B | 0.072 | BP, CC, and MF: all NS | 9.3-11.6% | 44.7% | 53.7-67.3% |
| EMX2 | 0.054 | **BP: 0.022** | 3.9-5.7% | 51.3% | 36.4-52.8% |
| LHX9 | 0.087 | BP: NS; **CC: 0.014** | 4.9-6.8% | 49.0% | 39.6-53.5% |
| LIN54 | 0.072 | **BP: 0.037; CC: 0.025** | 5.7-7.5% | 39.5% | 41.8-53.5% |
| MEOX2 | 0.042 | **BP: 0.015; CC: NS** | 3.6-5.7% | 50.5% | 34.9-52.2% |

### New targets of ARID3B, a gene linked to primary cilia sensory mechanism, carry out a conserved function in support of hypothesis H1A

ARID3B, a factor expressed in the Leydig cells of human and mouse testis (Samyesudhas, et al. 2014) that affects the expression of *Wnt1* and other genes in the placenta(Ali, et al. 2019), is highly and constitutively expressed throughout development in both turtle species and both sexes (Gessler, et al. 2023a). Here, ARID3B was more distant in regulome space between species than expected under the null hypothesis (H0) when applying trajectory analysis, and relatively low overlap was detected in the identity of ARID3B’s gene targets between species (Table 2). Yet, no significant differences in semantic similarity were detected when contrasting between versus within species comparisons, indicating a conservation of ARID3B’s function despite a change in gene targets (hypothesis H1A). This observation supports the notion that developmental systems drift affected the evolution of ARID3B’s gene targeting pattern between species. Of interest, when examining individual contrasts, semantic similarity was much lower between sexes in *Chrysemys* relative to within-*Apalone* contrasts, particularly for molecular function. Specifically, ARID3B targets at Cpi-FPT related to channel activity and membrane transport, which implicates ARID3B in the epigenetic regulation of TSD female development (Castelli, et al. 2020a; Weber, et al. 2020a), but mostly to cytoskeletal protein binding (tightly linked to primary cilia formation and maintenance) at Cpi-MPT. Notably, ARID3B terms relate directly to primary cilia for males incubated at 26°C in both species.

### EMX2, LHX9, LIN54, and MEOX2 are linked to primary cilia and support hypothesis H1B (same hub directs new function with different targets)

These TFs differed significantly more between *Chrysemys* and *Apalone* in their targeting patterns than expected under the null hypothesis H0 (Table 2), reflecting an evolutionary change in their regulatory subnetworks. Furthermore, they showed significant differences in functional annotation when testing for semantic similarity. EMX2 and MEOX2 showed a significant difference in semantic similarity for the biological process ontology, LHX9 for the cellular component ontology, LIN54 for both, and all showed changes in function that accompanied the change in gene targets between species (Table 2). The retention of the TF hub while gene targets changed in identity and function (hypothesis H1B) likely occurred via natural selection because the random nature of genetic drift is expected to result in a lack of overrepresented functions. Notably, all four TFs returned overrepresentation terms for primary cilia, while LIN54 is also related to Wnt signaling which primary cilia help sense(Long, et al. 2024). Additionally, all four TFs diverge somewhat among sex/temperature within-species networks (Table 2), revealing a lability between sexes or species that renders them candidates of interest underlying the transition of sex determination.

Interestingly, **EMX2**, a factor essential for gonadal and urogenital development eutherian mammals (Miyamoto, et al. 1997) and a marker of the bipotential gonad early in embryogenesis (Knarston, et al. 2020) when Wnt signaling also participates(Wilhelm, et al. 2025a), exhibited the lowest semantic similarity between the Asp-M31 network (an ancestral feminizing temperature) and all other *Apalone* networks. This result is consistent with previous genome-wide developmental-transcriptomic trajectory analysis showing the most sexually dimorphic gene expression in this GSD turtle occurs at the ancestrally feminizing 31°C temperature. Indeed, *Emx2* is upregulated in females at stages 19 and 22, and downregulated in males at 31°C, (Gessler, et al. 2023a), whereas in *Chrysemys, Emx2* shows monomorphic expression(Radhakrishnan, et al. 2017; Gessler, et al. 2023a). The opposite was not true, namely, the ancestral masculinizing temperatures (26°C) did not induce great divergence of the Asp-F26 network as no reduced semantic similarity was identified when comparing to all other *Apalone* networks. Consistent observations were made during the between-species comparisons, where the Asp-M31 networks returned fewer terms than all other networks, which overlapped largely with other *Apalone* networks, but not at all with *Chrysemys*, and not surprisingly, showed the lowest semantic similarity with *Chrysemys* networks. Terms in Asp-M31 were related to cell projection and microtubules – possibly pointing to general primary cilia related-terms (however explicit primary cilia terms were present in all other *Apalone* networks). Meanwhile, both *Chrysemys* networks returned terms related to anatomy and multicellular development with Cpi-FPT’s terms related to the nervous system and Cpi-MPT’s terms related to transcription and the immune system (functions linked to primary cilia in other systems as detailed in the discussion).

For **LHX9**, another bipotential gonad marker (Knarston, et al. 2020; Wilhelm, et al. 2025a) important for gonadal development in turtles and mammals(Birk, et al. 2000; Bieser and Wibbels 2014; Garcia-Alonso, et al. 2022; Lei, et al. 2022), the functional ontology terms of gene targets for *Apalone* were associated mostly with the cytoskeleton, which is key for the formation of primary cilia(May-Simera and Kelley 2012; Ge, et al. 2022). For *Chrysemys* terms were associated with ion channels many of which reside in the ciliary membrane. This supports LHX9’s mechanistically important role to relay temperature cues in TSD animals(Yatsu, et al. 2015; Czerwinski, et al. 2016), and suggests the evolution of GSD in *Apalone*’s lineage released LHX9 from its TSD function. Consistently, the within-species cellular component semantic similarity was generally quite high for *Apalone* and more moderate for *Chrysemys*, suggesting LHX9 may be more plastic in TSD *Chrysemys* and more canalized and perhaps regulating a more structural function in GSD *Apalone*. Consistently, *Lhx9* in *Chrysemys* is upregulated at MPT (26°C) throughout the thermosensitive period (stages 15, 19 and 22) whereas in *Apalone, Lhx9* retains this ancestral upregulation at 26°C throughout stages 15, 19, and 22, but it is also upregulated in stage 19 females compared to males irrespective of temperature, and in stage 22 females at 31°C(Gessler, et al. 2023a), revealing a clear evolutionary shift in expression and functional regulation with respect to sex, but not temperature.

**LIN54**, a factor involved in development and reproduction in *Drosophila*, *C. elegans* (Tabuchi, et al. 2011; Cheng, et al. 2017), and perhaps mammals (Tabuchi, et al. 2011; Hoareau, et al. 2024), is a core subunit of the DREAM/LINC complex that regulates DNA repair and the cell cycle(Schmit, et al. 2009; Tabuchi, et al. 2011; Marceau, et al. 2016; Bujarrabal-Dueso, et al. 2023; Hoareau, et al. 2024), a process dependent on the cytoskeleton and the centrosomes that comprise the primary cilia basal body, which become the mitotic spindle of dividing cells (May-Simera and Kelley 2012; Mill, et al. 2023). **LIN54** is expressed stably throughout development in both sexes in *Chrysemys* and *Apalone* (Gessler, et al. 2023a), yet it showed significant differences in semantic similarity for both biological process and cellular component between species. In *Apalone* these differences were largely driven by Asp-M26 which returned terms related to primary cilia, while other *Apalone* networks returned fewer or more general biological process terms, and cellular component terms related the nucleus. In contrast, *Chrysemys* terms were generally related to the nervous system and to neuron and ion/cation channel and cellular periphery at Cpi-FPT, but to the immune system, Golgi-vesicle transport, muscle cell-related, cytoskeleton/microtubule, nucleolus, and ribosome terms at Cpi-MPT. Importantly, previous studies have shown that many of these components have ties to primary cilia directly or indirectly [(Mill, et al. 2023) and references therein].

**MEOX2**, which is better known for its involvement in mesoderm differentiation (Candia, et al. 1992; Candia and Wright 1996) and limb development (Reijntjes, et al. 2007) but is also tied to nociception of inflammatory stimuli (Kokotović, et al. 2022) in vertebrates, is upregulated at FPT in *Chrysemys* at stage 12, while in *Apalone* its thermosensitivity changes between stages 12 (upregulated at 26°C) and 15 (upregulated at 31°C), and it is upregulated in stage 22 males developing at 31°C(Gessler, et al. 2023a). The biological process terms retrieved here for MEOX2 were strongly indicative of primary cilia and related components for *Apalone*, while *Chrysemys* networks were characterized by terms related to development for both networks with nervous system terms returned for Cpi-FPT and immune system for Cpi-MPT. Notably, although differences in the semantic similarity of cellular component were not significant for within versus between species comparisons, the semantic similarity value between male and female *Chrysemys* was very low (0.172), unlike in *Apalone* (> 0.5 for all contrasts), and due to the overrepresentation of calcium channel terms in Cpi-MPT but not Cpi-FPT networks, another important cellular component for relaying environmental cues during TSD gonadal development (Castelli, et al. 2020a; Weber, et al. 2020a), and consistent with recent work connecting calcium to male development via aldosterone production in *Trachemys* (Ye, et al. 2025).

### Targeted comparison of networks: subnetwork analysis also points to primary cilia

We also queried the networks qualitatively for subtler but potentially biologically important similarities and differences, focusing on hubs from highly supported subnetworks (those with edge scores Z > 10) and comparing (a) the identity of the TF hubs themselves, (b) the similarity in the identity of their gene targets, and (c) the functional annotations and overrepresentation of their gene targets. We identified 26 TFs of interest (Table 3) to further examine the molecular circuitry (network topology) of urogonadal development in *Chrysemys* and *Apalone*.

**Table 3.**
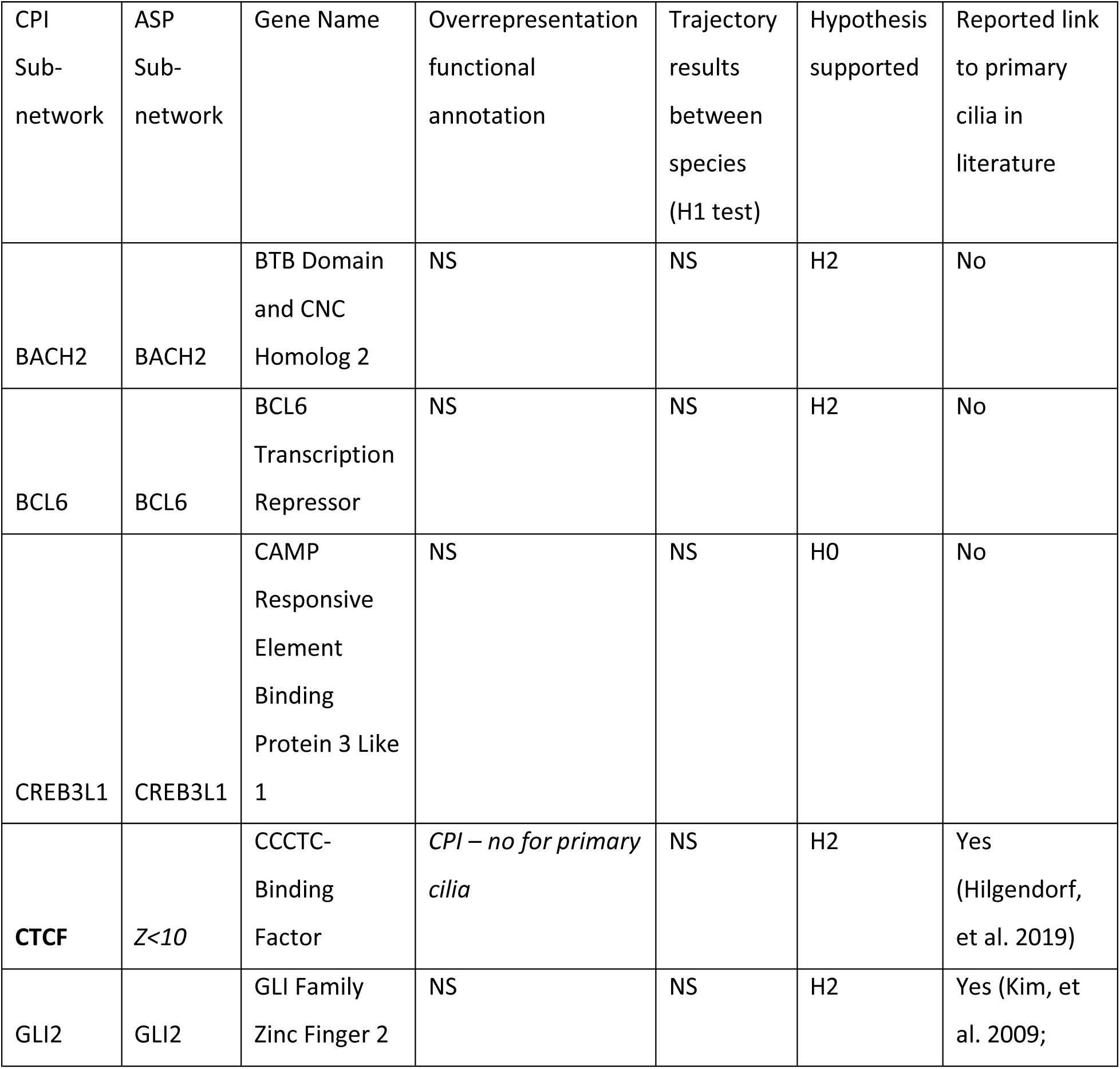
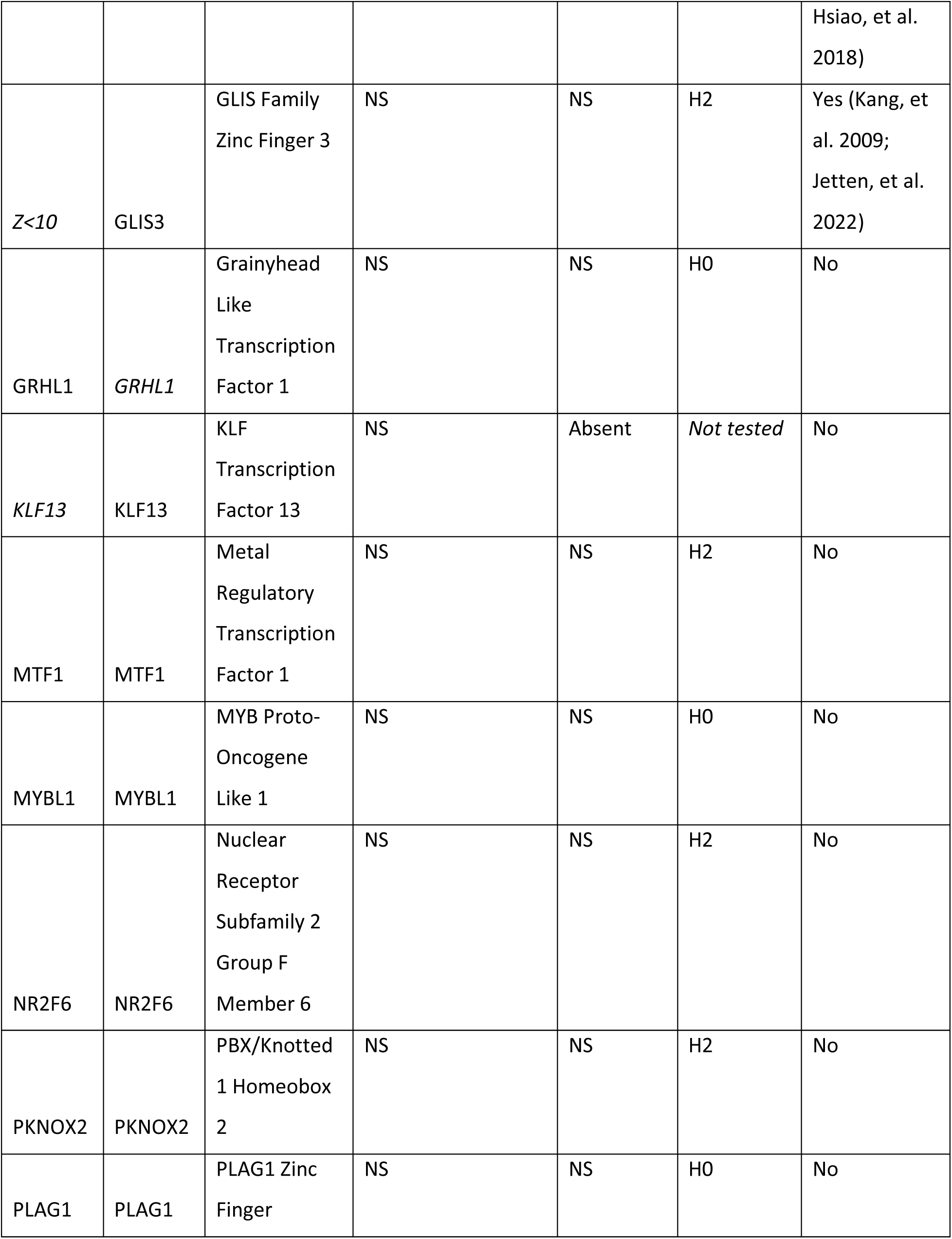
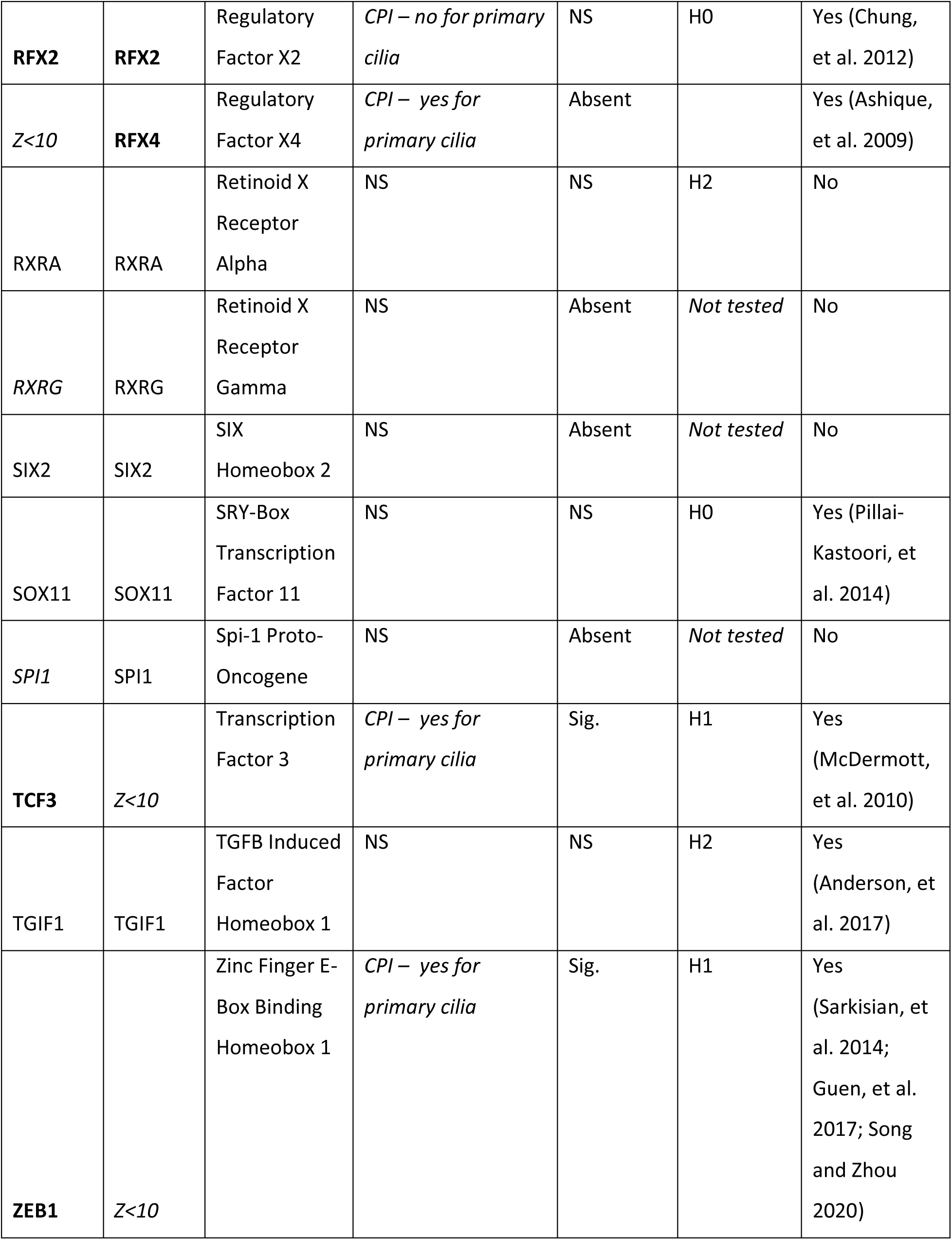
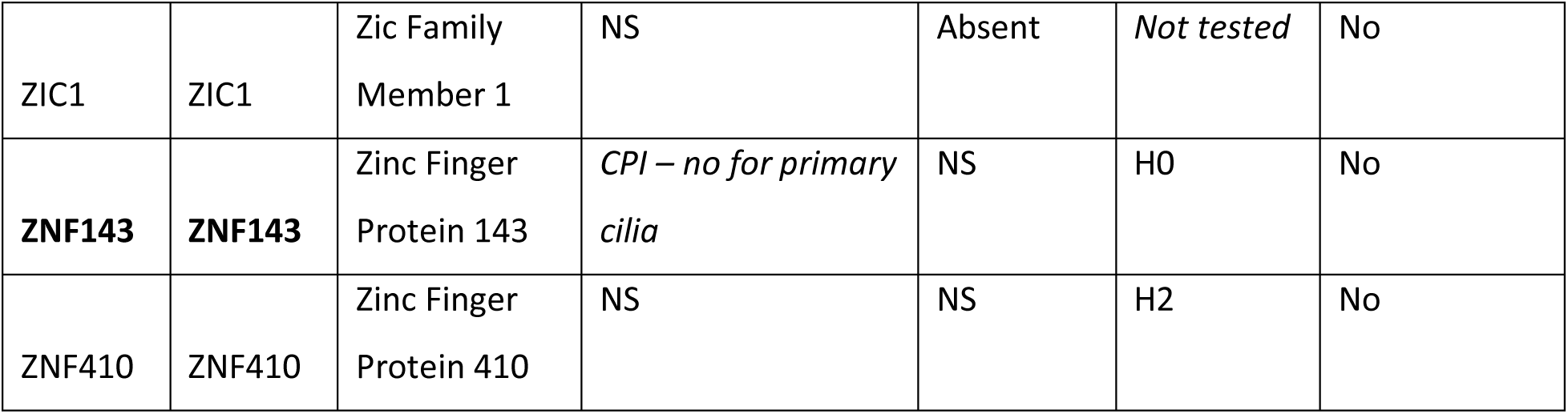
Highly supported subnetwork TF hubs identified in *Chrysemys* (CPI) or *Apalone* (ASP) using an edge weight cutoff of Z>10 (listed alphabetically). TFs that did not surpass the edge weight cutoff of Z>10 in one species are denoted as *Z<10*. *Italic font* denotes genes with low expression in one species [defined as log_2_(counts) < 1 for more than half of the libraries]. **Bold font** denotes hubs whose gene targets returned significant functional annotations in a species (detailed in Supplementary Materials). Significant trajectory results indicate the target genes diverged significantly between ortholog TFs. NS = nonsignificant; S = significant; Absent = excluded from trajectory analysis due to low expression; H0=fail to reject hypothesis H0; H1 = evidence supports H1; H2 = evidence supports H2.

The topology of these 26 TFs and their gene target interactions in the *Chrysemys* male and female subnetworks obtained from PANDA were nearly identical to each other in their pattern of gene targeting (Table 3), and the same was true within *Apalone* subnetworks. Thus, we focused on interspecific patterns. Between species, the same set of TF hubs were highly supported in both species, yet their highest supported gene targets differed considerably between *Chrysemys* and *Apalone*. Some TF hubs in the subnetwork were lowly expressed in only one species, maybe because they experienced a loss in activity (KLF13, RXRG, SPI1 in *Chrysemys*; GRHL1 in *Apalone*) yet they are returned as a hub likely because TF binding sites (and thus their regulatory potential) still exist in the promoter region of ancestral gene targets, maybe because they are still active in another context (e.g., pleiotropy).

Of these 26, six subnetwork hub TFs exhibited overrepresented biological functions exclusively in *Chrysemys* (CTCF, RFX2, RFX4, TCF3, ZEB1, and ZNF143) and virtually identical annotation terms at Cpi-MPT and Cpi-FPT for each TF, suggesting their likely role in general non-dimorphic sexual development. But importantly, gene targets for each of these six TFs differed greatly between species (0.68-11.5% overlap). This low overlap between species may be explained by (1) a functional loss for sexual development for these TFs in *Apalone* (hypothesis H1C, pseudofunctionalization), possibly due to genetic drift and consistent with an absence of significant overrepresentation results in this GSD turtle, or (2) a change in TF regulatory roles in *Apalone* compared to *Chrysemys*, suggesting the divergence in gene targets may have led to a change in how a particular functional process was regulated, perhaps by distributing the ancestral role across different TFs (subfunctionalization). Of note, three of these six TFs (TCF3, RFX4, and ZEB1) returned terms related to primary cilia. Moreover, RFX4 along with RFX2 play an important role in ciliogenesis, which is broadly conserved across vertebrates, and are testis regulators with RFX2 having a key role in spermatogenesis affecting ciliary and cytoskeleton remodeling genes (Vanwert, et al. 2008; Ashique, et al. 2009; Chung, et al. 2012; Kistler, et al. 2015; Wu, et al. 2016).

Furthermore, the terms related to primary cilia were overrepresented repeatedly irrespective of regulatory differences between species. Twenty TFs returned significant overrepresentation results related to primary cilia (ARID3B, EMX2, LHX9, LIN54, and MEOX2 mentioned above, plus DLX2, HIC2, HMBOX1, HOXD3, LHX2, MSX1, NFIA, NR2C2, RFX4, SHOX, SHOX2, SMAD4, SOX10, TCF3, and ZEB1), of which four have documented ties to gonadal development, i.e. EMX2 and LHX9 (described earlier), plus LHX2, MSX1, SMAD4, and SOX10 (Miyamoto, et al. 1997; Pellegrini, et al. 1997; Birk, et al. 2000; Mazaud, et al. 2002; Daftary and Taylor 2004; Itman and Loveland 2008; Polanco, et al. 2010; Le Bouffant, et al. 2011; Xie, et al. 2013; Sun, et al. 2016; Singh, Singh, Bhide, et al. 2022; Singh, et al. 2023). Of note, these terms tended to be present in Cpi-MPT and Asp-M26 networks, suggesting a bias towards male development tied to cooler temperatures.

### Shifts in regulation of known sexual development genes of interest

Taking a candidate gene approach, we also queried which TFs targeted several well-known gene regulators of sexual development: *Aromatase*, *Dhh, Dmrt1*, *Nr0b1 (Dax1)*, *Nr5a1 (Sf1)*, *Sox9*, and *Wt1*. The TFs among these genes of interest lacked position weight matrices in the JASPAR 2020 database used to obtain TF binding sites data but have been studied repeatedly in *Chrysemys* and *Apalone* (Valenzuela, et al. 2006; Valenzuela and Shikano 2007; Valenzuela 2008b, 2010; Valenzuela, et al. 2013; Radhakrishnan, et al. 2017; Mizoguchi and Valenzuela 2020; Gessler, et al. 2023a). We focused on expressed TFs targeting these genes of interest and identified those whose average edge weight difference between species-specific networks was greater than 3 (a difference of 3 between networks was chosen as qualitatively indicative of substantial differences because the top 5% of edges in these networks corresponded to values greater than 3). TBX20 emerged as a strong regulator of *Aromatase* in *Chrysemys*, and HNF4A, IRF1, and PAX3 as strong regulators of *Sox9* in *Apalone*, while *Dhh* was strongly targeted in *Chrysemys* compared to *Apalone* by NFIA and by CTCF, a gene affecting 3D chromatin structure which differs between turtles and other amniotes (Bista, et al. 2024) and is linked to male germline development(Rivero-Hinojosa, et al. 2021; Kitamura, et al. 2025). TFs that showed a greater targeting pattern of *Dhh* in *Apalone* relative to *Chrysemys* were ARID3B, EMX2, LHX9, and MEOX2, identified by our trajectory analysis described earlier, plus ALX1, BARHL2, BHLHE40, ELF5, HESX1, HEY2, HIF1A, ISL2, LHX2, LHX8, MSX1, MNT, and VAX1. Many of these TFs have previously been linked to sexual development and reproduction, with some related to gonadal establishment and development [EMX2 (Miyamoto, et al. 1997; Pellegrini, et al. 1997); LHX2 (Singh, Singh, Bhide, et al. 2022; Singh, et al. 2023); LHX9 (Birk, et al. 2000)]; ELF5 to epididymis (Bischof, et al. 2013; Wu, et al. 2013; Browne, et al. 2014); HIF1A to steroidogenesis in granulosa cells in the ovary (Baddela, et al. 2020); and others to germ cell development [BARHL2 to undifferentiated spermatogonia (Li, et al. 2024; He, et al. 2025); LHX8 to oocyte development (Choi, et al. 2008; Singh, Singh and Modi 2022); MSX1 to meiosis and germ cell migration (Le Bouffant, et al. 2011; Sun, et al. 2016)].

## Discussion

Extensive fragmentary data suggest the genetic architecture of vertebrate sexual development has evolved among disparate lineages at both the upstream regulators of sex determination and at the downstream mediators of sex differentiation, recycling some genes again and again (Crews and Bull 2009; Capel 2017; Stöck, et al. 2021). To our knowledge our study is the first to build and compare species-specific regulatory networks of gonadal development between closely related vertebrates with contrasting sex-determining mechanisms, using matched time-course sampling, to uncover putative key steps in the transitions in sex determination, by testing for conservation or divergence of modular components. Namely, we modeled gene-transcription factor interaction networks for two turtle species with and without sex chromosomes (*Apalone spinifera* and *Chrysemys picta*, respectively). TF hubs identified in *Chrysemys* were used as proxy for the ancestral TSD condition in turtles (Bista, et al. 2021) and compared to *Apalone*, a turtle with an evolutionarily derived ZZ/ZW GSD mechanism (Badenhorst, et al. 2013; Sabath, et al. 2016; Bista, et al. 2021), to assess the similarity of the identity and functional annotations of their gene targets. We note that these turtle lineages continued evolving since their split ∼175 Mya (timetree.org), such that not all differences between species may be attributable solely to changes in the softshell turtle family to which *Apalone* belongs. Indeed, further research with additional taxa is warranted to fully test the putative directionality of the evolutionary changes inferred here.

In general, gene regulatory networks evolve by altering hubs, their targets, or the strength of their connections, and we found evidence of all these processes at play during the evolution of turtle sex determination. Our results from the analysis of 89 TF hubs sufficiently expressed in the transcriptomes of *Chrysemys* and *Apalone* (Gessler, et al. 2023a) to enable testing, countered the hypothesis that the evolution of GSD required a complete overhaul of the regulatory network of sexual development (ruling out hypothesis H3, Fig 1). Instead, our findings revealed that most of these TFs (50 out of 89) support the null hypothesis H0 that some TFs hubs and their targets are conserved between species (likely representing core components of the regulatory network of turtle sexual development), while 39 other TFs diverged in the downstream genes they target (hypothesis H1). Some of these 39 TFs retained their ancestral function (hypothesis H1A), and some gained a new function (hypothesis H1B) (discussed below). Interpretation of H1C that TF targets lost their ancestral function is challenging as absence of evidence does not necessarily equate to evidence of absence because lack of annotations might reflect incomplete datasets instead. However, we did observe 15 of the 39 TFs returned significant functional annotations for one species but not for the other (considering all 5 ontologies tested against, which include molecular function, cellular component, biological process, PANTHER protein class, and PANTHER pathways). Among these, there was a strong bias (14/15 cases) towards absence of functional annotations in *Apalone*, so we interpret this result cautiously, as it would imply an extensive pseudofunctionalization of the molecular circuitry underlying sexual development in this GSD turtle that requires further functional validation. The one case which solely returned functional annotations for *Apalone* was ESRRG, a steroid receptor although most terms returned were general or related to RNA metabolism or gene expression. Interestingly, we found a single putative case of a TF (ZBED1) in the GSD *Apalone* that may have taken over the control of conserved targets (hypothesis H2) while also retaining its ancestral function, perhaps via developmental systems drift. Overall, our findings agree with the conservation of higher order regulatory network architecture documented in eukaryotes, and that substantial divergence has accrued in the identity and function of their regulatory targets, as observed across humans, flies, and worms (Boyle, et al. 2014). To date, few large comparative network studies exist, and more are needed to reveal common themes of network evolution (Karasawa and Koshikawa 2025). However, mechanistic studies have added important insights to knowledge of GRN patterns underlying evo-devo (Van Belleghem, et al. 2023) and our study contributes to this active field.

### Primary cilia underlie sexual development and key steps in the evolution of sex determination

Numerous lines of evidence from our results point to the important role that primary cilia play not only for TSD sexual development but also in the evolution of sex determination. First, five of the 39 TFs that changed targets between TSD and GSD turtles (EMX2, LHX9, LIN54, ARID3B, and MEOX2) could be functionally annotated across all six networks in turtles and showed significant results in the semantic similarity tests, enabling alternative hypotheses testing. All five returned overrepresentation terms for primary cilia, thus linking these organelles, previously documented as sensors of environmental and signaling pathway cues (Mill, et al. 2023), and involved in mammalian urogenital development (Wainwright, et al. 2014; Piprek, et al. 2019; Alves, et al. 2023), to evolutionary transitions in vertebrate sex determination explicitly for the first time, to our knowledge. Second, eight other TFs (HMBOX1, HOXD3, NFIA, NR2C2, SMAD4, SOX10, TCF3, ZEB1) that were significantly different in our trajectory analysis, but whose results were not comprehensive enough for hypothesis testing, also returned terms related to primary cilia. Third, several of the TFs whose regulatory functions may have been taken over at least partially by ZBED1 in *Apalone* are linked to primary cilia directly or indirectly (e.g. CTCF, GLI2, GLIS3, MTF1, NR2F6, PKNOX2, RXRA, TGIF1, ZNF410), supporting the notion that evolutionary shifts in the function of these organelles accompanied GSD evolution in this lineage. Fourth, ESRRG, the steroid receptor that returned functional annotations only for *Apalone,* regulates ciliary development (Wesselman, et al. 2023).

EMX2, LHX9, LIN54, and ARID3B have known links to sexual development and reproduction, and MEOX2 arises here as candidate for this new putative role. ARID3B likely evolved by developmental systems drift because its new target genes carry out the ancestral function (hypothesis H1A), while EMX2, LHX9, LIN54, and MEOX2 likely evolved by natural selection because their new targets in *Apalone* exhibit differences in their functional annotation from the putative ancestral targets in *Chrysemys* (hypothesis H1B). No evidence was found that the ancestral role of EMX2, LHX9, LIN54, and MEOX2 (i.e., the functions they orchestrate) might have been adopted by another TF (hypothesis H2). Only ARID3B returned primary cilia related terms in both species (Cpi-MPT and Asp-M26 networks), while the primary cilia terms for the other four TFs were found exclusively in *Apalone* networks: all four *Apalone* networks for EMX2 and MEOX2, while solely Asp-M26 for LHX9 and LIN54. This association with male development at cooler temperatures suggests that there may be important sex- or species-specific patterns associated with this structure, a hypothesis that requires future validation. We now discuss each of these five TF separately.

***Emx2*** is involved in the formation of cilia (Nguyen, et al. 2024) which help transduce Wnt signals important for gonadogenesis (Long, et al. 2024; Wilhelm, et al. 2025a), a process EMX2 also mediates in mammals (Miyamoto, et al. 1997; Knarston, et al. 2020). Our results show EMX2 was more divergent and returned fewer overrepresented terms for Asp*-*M31 (a network lacking strong primary cilia terms) than for all other networks (where primary cilia terms were strongly present), pointing to a potential reduced EMX2 functionality in Asp-M31, which we hypothesize prevents warmer temperatures from interfering with proper sexual differentiation of *Apalone* males at ancestrally-feminizing temperatures (Gessler, et al. 2023a). Our results may reflect a new role acquired in *Apalone* at late stages where *Emx2* differential transcription could induce differential ciliogenesis that might have evolved via natural selection, perhaps in concert with the evolution of *Meox2.* Indeed, we found evidence of regulatory coevolution for EMX2 and MEOX2. These two TFs clustered tightly within each species (Fig 2D) because their targets overlap more than any other TFs (by >70%) but also evolved substantially in parallel between *Chrysemys* and *Apalone*. This pattern could occur because they co-regulate the same genes, or because they compete for similar binding sites. This would be possible because TFs can overlap greatly in their active binding of targets yet induce different functional outcomes (via gene expression) due to the combinatorics of other aspects that fine tune regulation and influence the differential usage of shared binding sites (Holub, et al. 2024).

The observed coevolution of *Emx2* and *Meox2* is important because our study expands the putative roles for ***Meox2***, from mesoderm differentiation (Candia, et al. 1992; Candia and Wright 1996), limb development (Reijntjes, et al. 2007), interactions between placental epithelial and mesenchymal cells (Quinn, et al. 2000), and nociception (pain perception) of inflammatory stimuli (Kokotović, et al. 2022). *Meox2*’s thermosensitive and male-specific expression in turtles (Gessler, et al. 2023a) combined with its close association with EMX2, a known sexual development gene, render *Meox2* a novel candidate for a role in turtle sexual development whose evolution between *Chrysemys* and *Apalone* may have contributed to the transition in sex determination. Indeed, MEOX2 returned different biological process terms between turtle species, suggesting a role shift for this TF may have occurred. As this shift implicates the primary cilia, it could likely affect the thermosensory machinery, a notion further supported by the *Drosophila* homologue of MEOX2, *btn,* which mediates responses to noxious temperature (Kokotović, et al. 2022). Furthermore, within species, calcium channel terms were overrepresented for MEOX2 at Cpi-MPT but not Cpi-FPT networks, tying this TF to an important component of epigenetic regulation of TSD gonadal development (Castelli, et al. 2020a; Weber, et al. 2020a). And intriguingly, *Meox2* represses transcriptional co-activation by b-catenin (Kokotović, et al. 2022) a key ovarian development gene in TSD turtles (Mork and Capel 2013).

***Lhx9*,** another vertebrate gonadal development gene (Birk, et al. 2000; Bieser and Wibbels 2014; Knarston, et al. 2020; Garcia-Alonso, et al. 2022; Lei, et al. 2022; Wilhelm, et al. 2025a) appears to have shifted in the location and function of its protein targets between our focal TSD and GSD turtles, from channels and plasma membrane associated with calcium channels in *Chrysemys*, to nucleus, cytoplasm, and cytoskeleton but not calcium channel terms in *Apalone*. These changes are significant because calcium signaling plays an important role in TSD turtles like *Trachemys scripta* (Castelli, et al. 2020a), and the cytoskeleton affects primary cilia formation and maintenance (Ge, et al. 2022). Indeed, *Lhx9* is prone to developmental shifts despite its putatively critical role in the establishment of the gonad, because in *Trachemys*, *Lhx9* is expressed throughout the thermosensitive period but upregulated only in males at stage 15 (Czerwinski, et al. 2016), although it is also present in stage 26 ovaries (Barske and Capel 2010), while in *Chrysemys Lhx9* is upregulated in males throughout the thermosensitive period and in *Apalone* it is upregulated in females or in females developing at ancestrally feminizing temperature (Gessler, et al. 2023a). Such developmental systems drift affects other sexual development genes across turtles and vertebrates (Valenzuela, et al. 2013; Ge, et al. 2017; Mizoguchi and Valenzuela 2020).

***Lin54*,** a core subunit of the DREAM/LINC complex which is highly conserved across animals and plants (Schmit, et al. 2009; Tabuchi, et al. 2011; Marceau, et al. 2016; Bujarrabal-Dueso, et al. 2023; Hoareau, et al. 2024) functions as activator and repressor by targeting different gene sets (Tabuchi, et al. 2011) and plays roles in development, reproduction in invertebrates (Tabuchi, et al. 2011; Cheng, et al. 2017) and perhaps mammals (Tabuchi, et al. 2011) where a paralog of *Lin54*, *Mtl5*, has testis-specific action during spermatocyte meiosis(Hoareau, et al. 2024). Interestingly, LIN54 favors binding to autosomes in the soma and influences the X chromosome gene expression indirectly in *C. elegans* (Tabuchi, et al. 2011), where it helps DNA repair (Bujarrabal-Dueso, et al. 2023). In turtles, we observed significantly overrepresented terms related to development for LIN54 targets in *Chrysemys*, and to primary cilia in *Apalone.* But because LIN54 is related to Wnt signaling (sensed with the help of primary cilia), which is important for gonadogenesis in TSD and GSD turtles (Rhen, et al. 2021; Zhou, et al. 2023) and other vertebrates (Capel 2017), our results render *Lin54* an interesting candidate ever-present and primed to help transduce differential Wnt signaling during sexual development.

***Arid3b*** is expressed in numerous cancer types including breast and ovarian cancer (Oguz Erdogan, et al. 2014; Samyesudhas, et al. 2014; Bobbs, et al. 2015; Saadat, et al. 2021) perhaps because it regulates the cell cycle and stem cell genes (Saadat, et al. 2021) as it is a member of the LIN28-let-7-ARID3B pathway that promotes cell proliferation (Ali, et al. 2019). While not itself a member of the DREAM complex, ARID3B binds to E2F and RB family genes (Saadat, et al. 2021) which are important members of the DREAM complex (Hoareau, et al. 2024), and along with LIN54, regulate the mitotic gene *Cdc2 (Schmit, et al. 2009; Saadat, et al. 2021)*. ARID3B is also expressed in the Leydig cells (Samyesudhas, et al. 2014), and binds to *Wnt1* (Ali, et al. 2019). Importantly, our results showed that ARID3B was most differential between *Chrysemys* networks, where it targets genes related to channel activity and membrane transport at Cpi*-*FPT in agreement with the reported regulation of TSD female development via the phosphorylation of STAT3 mediated by calcium channels, which represses *Kdm6b* expression and consequently, the epigenetic activation of *Dmrt1* and downstream male-differentiation genes (Weber, et al. 2020a). Our observations render ARID3B an important upstream candidate for male and female development in *Chrysemys* with putative opposite action to MEOX2 which exhibited overrepresentation of calcium channel terms at Cpi-MPT but not Cpi-FPT networks in this TSD turtle.

### Evolution of *Dhh*, a known candidate gene, is also linked to the turnover in sex determination associated to primary cilia

We previously identified *Dhh* (Desert Hedgehog Signaling Molecule) as a gene with sex-specific expression that switched between *Chrysemys* and *Apalone* (Gessler, et al. 2023a). *Dhh* is upregulated at FPT (31°C) during the thermosensitive period in *Chrysemys*, and in males or at 26°C in *Apalone* at the same stages 15-22 (Gessler, et al. 2023a). In contrast, *Dhh* was not expressed during a similar time window in *Trachemys scripta elegans* (stages 15 – 19 and 21) (Czerwinski, et al. 2016). *Dhh* helps mammalian testis development and is upregulated in male mice during stages e11.6-e12.0 that correspond to turtle stages 18-21 (Munger, et al. 2013; Czerwinski, et al. 2016; Pachernegg, et al. 2022). *Apalone’s* transcriptional pattern agrees with mammalian *Dhh* expression and its role in testis development, suggesting an evolutionary shift between *Apalone* and *Chrysemys* lineages in *Dhh* regulation. Our results show that the TFs NFIA and CTCF increased their targeting of *Dhh* in *Chrysemys* relative to *Apalone*, and 17 other TFs have increased targeting of *Dhh* in *Apalone* relative to *Chrysemys,* including LHX9, MEOX2, EMX2, and ARID3B related to primary cilia, plus BARHL2, LHX2, ISL2, ALX1, HESX1, LHX8, ELF5, HIF1A, HEY2, MNT, VAX1, BHLHE40, and MSX1 [implicated in female pathways via active male downregulation (Capel 2017)].

Hedgehog signaling is dependent on primary cilia in vertebrates, particularly for hedgehog genes *Shh* and *Ihh* (Bangs and Anderson 2017), but also *Dhh (Nygaard, Almstrup, Lindbæk, et al. 2015)*. Specifically, hedgehog signaling components were found in primary cilia of immature Leydig cells (Yao, et al. 2002; Nygaard, Almstrup, Lindbæk, et al. 2015), whose differentiation is induced by DHH signaling, thus rendering DHH signaling via primary cilia a potential regulator of Leydig cell recruitment and or differentiation (Yao, et al. 2002; Barsoum, et al. 2009; Nygaard, Almstrup, Lindbæk, et al. 2015). DHH signaling is also present in mouse ovary shortly after birth where it participates in theca cell differentiation (Liu, et al. 2015), pointing to a general role in specification of steroidogenic cells in the gonad. Furthermore, a direct link between DHH signaling and primary cilia via a Type II non-canonical cilia signaling mechanism was detected in the developing mouse heart (Fulmer, et al. 2020).

### Identification of primary cilia as organelles mediating sex determination extends the calcium and redox (CaRe) model into a novel *Primary Cilia Integration* hypothesis

Primary cilia are antennae-like organelles, now recognized as essential for the perception and transduction of signals in most cell types, whose disfunction is linked to numerous diseases, including hypogonadism and genitourinary disorders of development [reviewed in (Mill, et al. 2023)]. The primary cilium consists of a basal body made up of the centriole (which moonlights as the mitotic spindle), a transition zone (the cilium gateway), and an axoneme typically made up of nine microtubule doublets (Mill, et al. 2023; Long, et al. 2024). Proteins are moved up and down the axoneme via anterograde (IFT-A, kinesin) and retrograde (IFT-B, dynein) transport. The ciliary membrane can contain numerous proteins including TRP channels, which are important for relaying Ca^2+^, to communicate environmental changes such as temperature, mechanical force, and other signals, and is key to the primary cilia’s ability to integrate environmental inputs to the cell (Decaen, et al. 2013; Delling, et al. 2013; Nauli, et al. 2016). Primary cilia are also involved in relaying numerous signaling pathways beyond hedgehog and Wnt, including GPCR, TGF-B/BMP, NF-kB, among numerous others (Wann, et al. 2014; Bangs and Anderson 2017; Anvarian, et al. 2019; Jiang, et al. 2019; Lee 2020; Mill, et al. 2023). Thus, primary cilia are uniquely suited to integrate environmental cues into developmental outputs, such as is essential for developmental plasticity.

Our findings suggest a link between primary cilia and turtle sexual development, expanding previous reports showing they play a critical role in the development of the urogenital ridge (Wainwright, et al. 2014), Wolffian ducts (mediated by DHH signaling) (Alves, et al. 2023), and somatic and germline gonadal components in mammals (Wainwright, et al. 2014; Piprek, et al. 2019) and perhaps also in *Paralichthys olivaceus*, a fish with thermosensitive XY system (Wang, et al. 2020). Namely, TFs with overrepresented terms related to the primary cilia present in *Chrysemys* and *Apalone*, include ARID3B, EMX2, LHX9, LIN54, and MEOX2 described above, plus DLX2, HIC2, HMBOX1, HOXD3, LHX2, MSX1, NFIA, NR2C2, SHOX, SHOX2, SMAD4, SOX10, and ZEB1,of which LHX2 (Singh, Singh, Bhide, et al. 2022; Singh, et al. 2023), MSX1 (Le Bouffant, et al. 2011; Xie, et al. 2013; Sun, et al. 2016), SMAD4 (Itman and Loveland 2008), and SOX10 (Polanco, et al. 2010) have previously known ties to gonadal development. Because primary cilia are organelles found in nearly all vertebrate cell types that help cells understand the context of cues from the environment or signaling pathways, including Wnt signaling (Lee and Gleeson 2010) and hedgehog signaling (Bangs and Anderson 2017), their biology integrates and expands upon the calcium and redox (CaRe) hypothesis of sex determination which proposes that thermosensitive cytoplasmic calcium and mitochondrial redox signaling interactions activate or repress male- and female-specific developmental pathways (Castelli, et al. 2020a), as primary cilia are linked to both reactive oxygen species (ROS) and calcium signaling.

Thus, we propose an expanded model of sex determination to incorporate the role of primary cilia – the *Primary Cilia Integration* hypothesis (Fig 3). We hypothesize primary cilia to be the antennae relaying environmental cues and integrating them via the signaling pathways they mediate (Decaen, et al. 2013; Delling, et al. 2013; Nauli, et al. 2016; Mill, et al. 2023) to guide sex determination and differentiation in TSD turtles. Warmer temperatures mediate calcium signaling through TRP channels present in ciliary membranes (Köttgen, et al. 2008; Decaen, et al. 2013; Delling, et al. 2013) and TRP genes are implicated in TSD biology (Yatsu, et al. 2015; Castelli, et al. 2020a; Weber, et al. 2020a). Calcium fluxes through TRP channels that open at warmer temperature induce phosphorylation of STAT3 which inhibits *Kdm6b* expression, thus favoring female developmental pathways (Ge, et al. 2017; Weber, et al. 2020a; Wu, et al. 2024). Additionally primary cilia transduce Wnt signaling (May-Simera and Kelley 2012; Oh and Katsanis 2013; Lee 2020), hedgehog signaling (Nygaard, Almstrup, Lindbæk, et al. 2015; Fulmer, et al. 2020; Alves, et al. 2023), NFkB signaling (Wann, et al. 2014), and are negatively regulated by NRF2 signaling (Liu, et al. 2020; Morleo, et al. 2023), all pathways with known or suspected links to sex determination (Castelli, et al. 2020a). Wnt’s and hedgehog’s signaling role in gonadal development is well documented: (1) Wnt canonical signaling is linked to female development (Yao 2005; Mork and Capel 2013; Rhen, et al. 2021; Wilhelm, et al. 2025a), increases under female producing temperatures in the snapping turtle *Chelydra serpentina* (Rhen, et al. 2021), and failure to inhibit it in humans disrupts male development (Lundgaard Riis, et al. 2024); (2) hedgehog canonical signaling is linked to male development and is entirely dependent on the primary cilium (Yao, et al. 2002; Nygaard, Almstrup, Lindbæk, et al. 2015; Bangs and Anderson 2017; Alves, et al. 2023); and (3) Wnt and hedgehog signaling can be mutually antagonistic (Ding and Wang 2017) consistent with the mutual inhibition required for alterative commitment to male or female developmental fate (Capel 2017).

**Fig 3.**
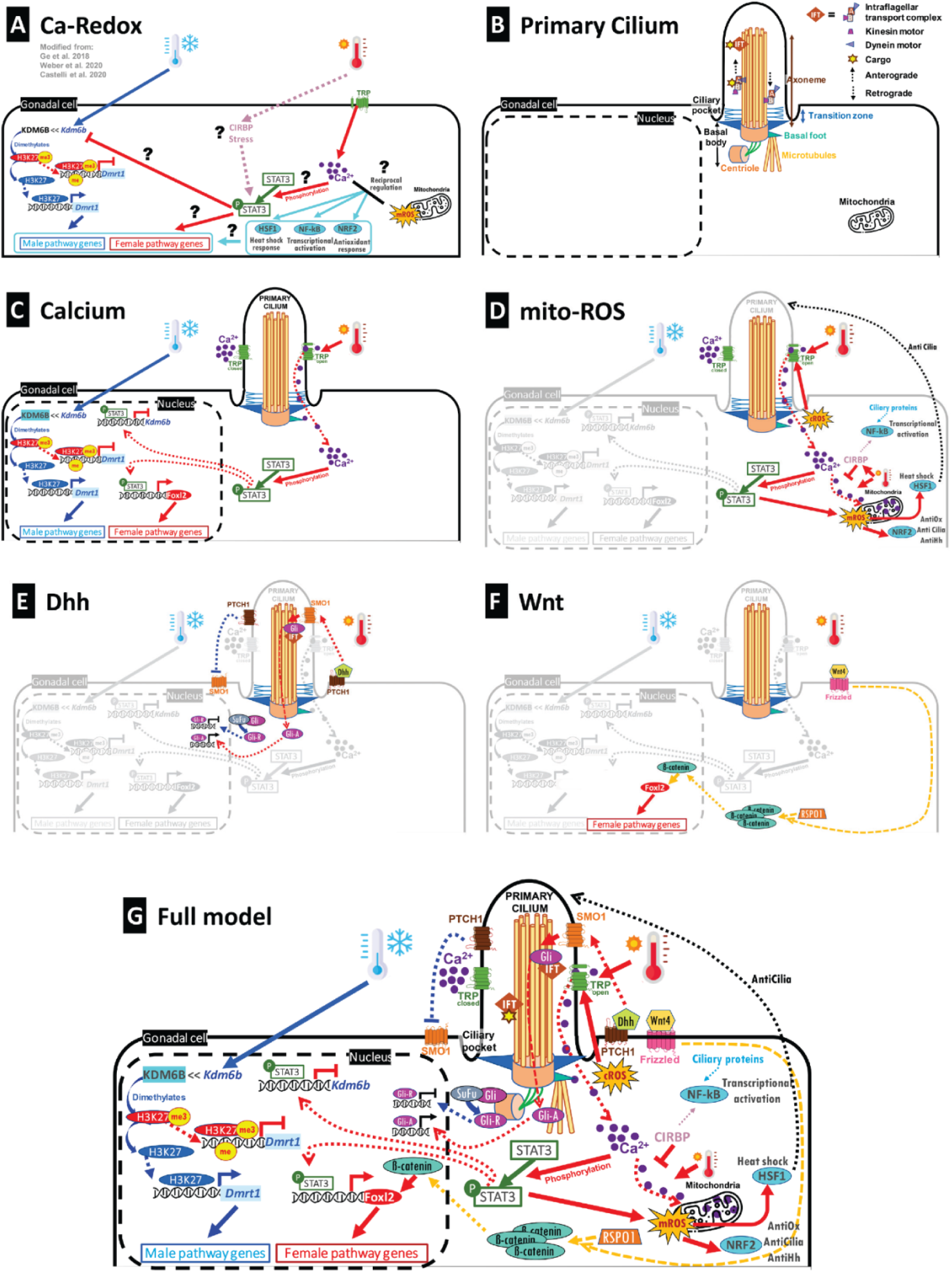
*Primary cilia integration* model of temperature-dependent sex determination and differentiation. The model is informed by findings in this study from *Chrysemys picta* and *Apalone spinifera* and extended from the literature on other TSD turtles and GSD vertebrates (see text for further details). A) Calcium-Redox model where thermosensitive calcium-redox signaling activates or represses sexual development pathways. B) Primary cilium structure. C) Calcium signaling mediated through the primary cilium with hypothesized effect on molecular sex determination mechanisms. D) Calcium signaling mediated through the primary cilium with hypothesized cross talk to mitochondrial reactive oxygen species and corresponding pathway activation. E) Thermosensitive hedgehog signaling mediated through the primary cilium. F) Thermosensitive Wnt signaling potentially mediated by the primary cilium with hypothesized influence on sex determination pathways. G) Full *Primary Cilia Integration* model. TRP channels open at warm temperatures, allowing Ca^2+^ influx into the primary cilium. This calcium can be relayed to the cell and transported to the mitochondria, contributing to the production of reactive oxygen species (ROS), altering the CaRe status of the cell. At lower temperatures, TRP channels are closed, blocking entry of Ca^2+^. CIRBP, which inhibits movement of Ca^2+^ into the mitochondria, may help modulate the CaRe state. CaRe crosstalk may influence signaling pathways like HSF1, NF-kB, and NRF2 (Castelli, et al. 2020b), which have known ties to the primary cilium (Martin-Hurtado, et al. 2019; Mc Fie, et al. 2020), of which NF-kB signaling is mediated by CIRBP (Mc Fie, et al. 2020). When high temperatures raise Ca^2+^ levels, STAT3 is phosphorylated by a still unknown factor. pSTAT3 inhibits transcription of *Kdm6b*, a histone demethylase (Weber, et al. 2020b). Failure to produce its protein results in retention of silencing chromatin marks (H3K27me3) at the *Dmrt1* promoter, inhibiting expression of male pathway genes. Furthermore, pSTAT3 binds to the *Foxl2* promoter, driving downstream expression of female pathway genes (Wu, et al. 2024). *Foxl2* expression may be further enhanced by the accumulation of B-catenin in response to Wnt signaling which is stabilized by RSPO1 (Wilhelm, et al. 2025b). Wnt signaling can be mediated by the primary cilium (Mill, et al. 2023) and is thermosensitive in TSD turtles (Shoemaker, et al. 2007). When STAT3 is unphosphorylated at cooler temperatures, *Kdm6b* is expressed and can activate *Dmrt1* through demethylation of H3K27me3, driving expression of male pathway genes. DHH, another signaling pathway important for sexual development and dependent on the primary cilium (Nygaard, Almstrup, Lindbaek, et al. 2015), may be temperature sensitive and evolutionarily labile (Gessler, et al. 2023b). When activated, DHH ligands binds to PTCH1 receptors which allows activation of SMO. SMO converts GLI transcription factors into active forms that can activate downstream target genes.

This expanded model reveals several new questions to guide future studies in turtles, and multiple aspects of the model must be functionally validated, such as exploring the thermosensitivity of hedgehog, identifying the agent which phosphorylates STAT3, and characterizing the makeup of ciliary membranes. The evolutionary changes revealed by our analyses as described earlier suggest that important modifications might have occurred at several levels in *Apalone* compared to *Chrysemys*, including in the rate of ciliogenesis, in the morphology and composition of the primary cilia, and in the location of ciliary proteins (ciliary versus extraciliary) which can affect their functions (altering cell cycle regulation, cytoskeletal regulation, and trafficking) (Hua and Ferland 2018). Additionally, the species-specific role of each of the candidate TFs identified here must be tested. Another important question is if, or how, signals from sex chromosomes may interface with the primary cilia as would be predicted if primary cilia underlie the evolution of vertebrate sex determination? For instance, *Wt1*, the Wilm’s tumor protein 1 gene involved in the development of the bipotential gonad and later testes (Gao, et al. 2006) or ovaries depending on the spliceoform (Gregoire, et al. 2023), causes Wilm’s tumors which are associated with primary cilia disfunction (Kitamura, et al. 2020), and intriguingly, *Wt1* is linked to the sex chromosomes in *Glyptemys insculpta* and *Siebenrockiella crassicollis* turtles, two species with independently evolved XY systems (Montiel, et al. 2016). Likewise, *Dmrt1*, a testis development gene tied to testicular germ cell cancer which is also associated with primary cilia disfunction (Litchfield, et al. 2016), is linked to the sex chromosomes of *Staurotypus triporcatus* turtles, another lineage with an independently evolved XY system (Montiel, et al. 2016). Further, the steroidogenic factor 1 gene *Sf1*, a target and partner of *Wt1* during gonadal development (Wilhelm, et al. 2025b), is tied to metabolic homeostasis (Dhillon, et al. 2006) that primary cilia mediate (Yang, et al. 2021), and is linked to the ZW sex chromosomes of *Apalone spinifera* softshell turtles (Lee, et al. 2019). Emerging resources, such as the development of turtle organoids (Zdyrski, et al. 2024), will provide an excellent functional genomics resource in which to investigate the potential role of primary cilia in sensing the environment during sex determination and differentiation for TSD species. Indeed, previous studies have examined the effects of primary cilia in mammary organoids (McDermott, et al. 2010; Guen, et al. 2017).

## Conclusions

We generated gene-transcription factor sexual development networks for two turtle species, *Chrysemys picta* and *Apalone spinifera*. While generalized in scope due to sample pooling, the characteristics of these networks are consistent with previously reported biology of sexual development. These networks represent mechanistic models to inform future investigations into the evolution of sex determination in turtles and vertebrates. Our results support the following conclusions: (1) There is a large degree of conservation in transcription factor hubs between *Chrysemys* and *Apalone*, consistent with the prevailing understanding of a high degree of conservation in elements of vertebrate sex determination networks and evo-devo toolkit hypotheses. (2) While the TF hubs were conserved, the targets of those hubs often were not. In some cases, these varying targets converged on similar functional annotations, suggesting a role for developmental systems drift. In other cases, they significantly differed in their functional annotations suggesting a possibility of natural selection or genetic drift to be at play. (3) We identified one TF with conserved ancestral targets that may have undergone developmental systems drift through acquisition of additional gene targets during GSD evolution. (4) We identified several candidate TFs in our analysis that are of interest for sex determination that have known roles in mammalian sexual development and expand this role to reptiles. (5) We identified predicted regulatory changes in *Dhh* that could underpin its male-to-female shift in gene expression previously observed between *Apalone* and *Chrysemys*. (6) Finally, we found primary cilia to be tightly linked to sex determination in both TSD and GSD species and uncovered a potential role for primary cilia as the sentinels of environmental signaling in TSD, explicitly linking them to this function while expanding upon the CaRe Hypothesis, and tying them to transitions in sex determination for the first time. Given the ability of the primary cilium to interface with the environment and with so many signaling pathways (several with known ties to sexual development), it is tempting to hypothesize that they could underly the evolutionary diversity observed in vertebrate sex determination.

Further research is warranted to test these hypotheses, and should include comparative proteomics of primary cilia in developing gonads of TSD and GSD species to elucidate finer details of their compositional dynamics during cell differentiation at various embryonic stages of sexual development. This approach is still missing both in our general understanding of primary cilia function and in their role in development and disease (Mill, et al. 2023), but will also help decipher the role of primary cilia in the molecular and cellular evolution of plasticity and canalization.

## Materials and Methods

We constructed models of gene-TF interaction networks for both *Chrysemys* and *Apalone* using PANDA as implemented in the R package pandaR (Glass, et al. 2013; Schlauch, et al. 2017). As input to construct the networks, pandaR takes gene expression data, transcription factor binding site data, and protein-protein interaction data. PANDA utilizes a message passing algorithm to integrate and find agreement among multiple datasets, *via* similarity calculations, building a consensus network with predicted regulatory relationships (Glass, et al. 2013). We describe the generation of each of those datasets below.

### Gene expression dataset

We used RNA-seq data from a previous study (Gessler, et al. 2023a), which we generated by incubating eggs from both *Chrysemys* and *Apalone* turtles at temperatures that are either 100% male- or female-producing in *Chrysemys* (MPT: 26°C and FPT: 31°C, respectively) following our standard protocols(Valenzuela 2009). These temperatures are within the optimal developmental range for both species and can be used as a proxy for the ancestral TSD condition for trionychid softshell turtles such as *Apalone* whose GSD sex determination mechanism is evolutionarily derived (Bista, et al. 2021). This is because the TSD mechanism in the closest relative of *Apalone*’s family, *Carettochelys insculpta*, is likely not ancestral but secondarily derived, the result of a putative GSD-to-TSD reversal (Valenzuela and Adams 2011b; Literman, Burret, et al. 2018), and *Apalone* is equally related to *Chrysemys* as to all other TSD turtles in the suborder Cryptodira. Tissue was collected from both species across five matched stages of development [sensu (Yntema 1968)] and included the following tissues: stage 9 – trunks, stages 12 and 15 – adrenal kidney gonad complexes (AKGs), and stages 19 and 22 – gonads. Stages 9 and 12 precede the thermosensitive sex determination period (TSP) in *Chrysemys*, stage 15 sits at the onset of the TSP, whereas stages 19 and 22 are at the mid and late TSP. Embryos of *Chrysemys* were presumed to be developing males or females according to their incubation temperature, while *Apalone* embryos were sexed by PCR using molecular markers (Literman, et al. 2017), an improved sexing technique compared to qPCR of rRNA genes (Literman, et al. 2014). Thus, *Apalone* samples correspond to a full factorial dataset with male and female embryos developing under temperatures that represent the ancestral MPT and ancestral FPT. Two biological replicates (RNA libraries) were generated for each condition (species by stage by temperature, and by sex for *Apalone*), and each library included RNA pooled from 11-15 embryos. At least 40 M clean Illumina 150 bp paired-end reads were generated per library (with a 94%–97% retention rate per library). Further details of the transcriptomic datasets were previously reported (Gessler, et al. 2023a). Reads were trimmed with Trimmomatic (v0.36) (Bolger, et al. 2014) and mapped with GSNAP (v20170317) (Wu and Watanabe 2005; Wu and Nacu 2010) to their respective reference genome: *Chrysemys* GCF_000241765.3_Chrysemys_picta_bellii-3.0.3 (Badenhorst, et al. 2015) and *Apalone* BioProject: PRJNA837702(Bista, et al. 2024). Using StringTie (v1.3.4)(Pertea, et al. 2015), reads were assembled into transcripts by library, merged, and their abundance was calculated. Then, transcripts and counts were consolidated into gene models with tximport (v1.10.1) (Soneson, et al. 2015). Finally, counts were TMM-normalized to correct for library size and log_2_-transformed to correct for heteroskedasticity using EdgeR (v3.24.3) (Robinson, et al. 2010).

### Transcription factor binding site (TFBS) motif dataset

Promoter sequences, defined as -1.5Kb and +500bp surrounding the transcription start site, were extracted for all annotated genes from the *Chrysemys* and *Apalone* reference genomes. Because DNA binding domains for transcription factors are highly conserved in vertebrates (Lowry and Atchley 2000; Schmidt, et al. 2010; Nitta, et al. 2015), we used vertebrate transcription factor binding matrices from the open access JASPAR Core Vertebrate database (v2020) (Fornes, et al. 2019). TF position weight matrices were searched against these promoter sequences, using CiiiDER (v1.10.6) (Gearing, et al. 2019) with the deficit parameter set to 0.05 to create a map indicating whether promoters contained putative binding sites for these transcription factors.

### Protein interaction dataset

Protein sequences and protein-protein interaction data from vertebrates were obtained from the STRING database (v11.0) (Szklarczyk, et al. 2019) which included data from 41 vertebrates (30 mammals, 3 birds/reptiles, 1 amphibian, and 7 fishes). In parallel, we obtained the longest translated CDS for proteins present in the *Chrysemys* genome (Badenhorst, et al. 2015). Because PANDA focuses on gene-transcription factor interactions, the list of *Chrysemys* proteins was filtered down to retain transcription factors also present in the JASPAR core vertebrate database (v2020). Using a reciprocal best blast approach (blastp), we compared the sequences of the STRING interacting proteins to the subset of *Chrysemys* transcription factors and did a final filtering step resulting in a list of TFs with a percent identity of ≥90% and resulting query coverage of ≥99%. STRING proteins and their interactors that passed through these filters were retained and assumed to represent protein interactions likely present in *Chrysemys* due to their high level of homology. For *Apalone*, protein sequences from *Chrysemys* representing putative interacting proteins were searched (tblastn) against the *Apalone* genome (Bista, et al. 2024) and their presence was confirmed, such that the same protein interaction file was used as input for both species.

### Building models of gene – transcription factor interaction networks with PANDA

For *Chrysemys*, each gene expression dataset per temperature (26°C and 31°C) consisted of ten gene expression libraries, encompassing two biological replicates from five developmental stages, yielding 20 libraries total. These temperature-specific libraries were used as input to predict MPT- and FPT-networks in PANDA (Cpi-Female-31°C network and Cpi-Male-26°C network). Because each incubation temperature produced both sexes in *Apalone*, there were 40 datasets for this GSD species, i.e. two replicates per stage for each sex-by-temperature combination. Sex-by-temperature sets were used as input in PANDA to generate 4 networks (Asp-Female-26°C, Asp-Female-31°C, Asp-Male-26°C, and Asp-Male-31°C). We will refer to these networks as Cpi-FPT, Cpi-MPT, Asp-F26, Asp-M26, Asp-F31, and Asp-M31 hereafter.

The PANDA algorithm was implemented with the R package pandaR (v1.22.0) (Schlauch, et al. 2017), using as input the gene expression, TFBS motif, and protein-protein interaction datasets described above. Default settings were used in most cases, but the mode was set to ‘intersection’ to only include TFs present in the gene expression dataset while keeping network size to a manageable scale. Following network construction, it was discovered that the mode option works differently than described in the pandaR manual, so we manually confirmed which TFs in the final network were expressed in our transcriptomes and disregarded unexpressed TFs when interpreting the results. Networks produced by PANDA consist of gene-by-TF matrices populated with similarity scores (akin to Z scores) describing the likelihood that an edge (i.e. the connection between a gene and TF that defines a regulatory relationship) is true, where positive values represent greater support for an edge and negative values represent lesser likelihood that an edge is true.

To test for differences between networks, we used a permutation approach implemented with a custom R script to build a test distribution of networks (Fig 4). For this, we first built a gene (row) by library (column) matrix populated with gene expression values (20 columns in the case of *Chrysemys*). Then, we ran 999 permutations randomizing the order of the libraries (columns) in the matrix. The resulting 999 permutated matrices were split in half (columns 1-10, columns 11-20 for *Chrysemys*) to generate two sets of 999 matrices with permuted columns, and a network was generated (as described above) from each set to obtain two random distributions of ‘permuted’ networks against which to test each empirical network. The empirical FPT network was tested against one distribution while the empirical MPT network was tested against the second distribution. Likewise, for *Apalone*, the initial 40-column matrix (for the 40 libraries) was also permuted 999 times, and these randomized matrices were split in four groups (columns 1-10, 11-20, 21-30, 31-40) to generate four sets of 999 permuted matrices. A network was generated from each set of permuted matrices in PANDA to obtain four random distributions against which to test each empirical network (Asp-F26, Asp-M26, Asp-F31, and Asp-M31).

**Fig 4.**
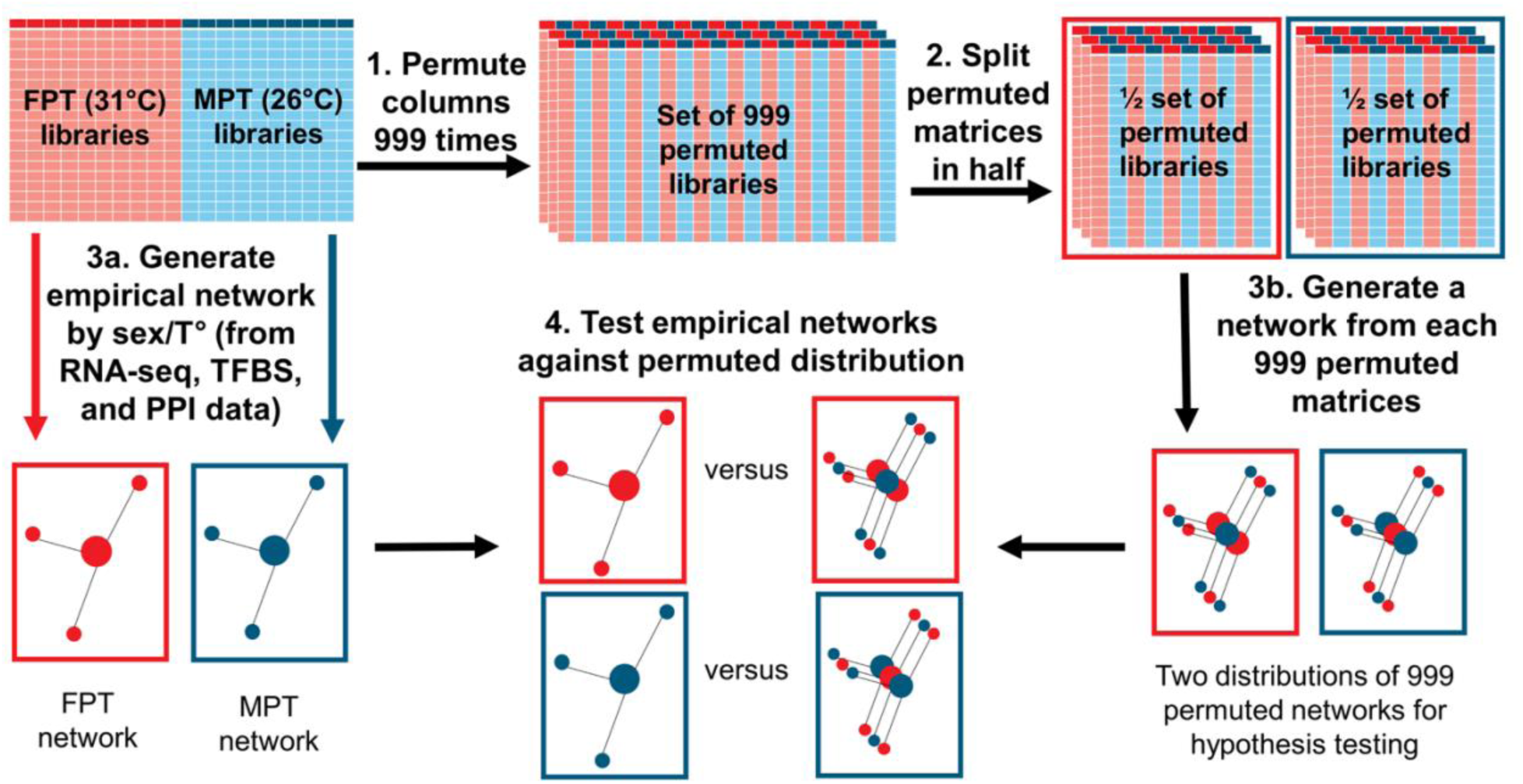
Pipeline of the permutation procedure used to generate random distributions for network comparisons using the *Chrysemys* data as example. First, a matrix containing normalized gene expression values for all 20 libraries (red columns = FPT/female libraries, blue columns = MPT/male libraries) were permuted 999 times to generate 999 matrices with random column order. Second, each of these 999 matrices were split in half. Third, each 10-column set of submatrices was used as input to generate a random distribution of networks. Fourth, each empirical network derived from the original libraries was tested against one of the random distributions. The same process was used for *Apalone*’s 40 libraries for *Apalone* as detailed in the text. TFBS=Transcription Factor Binding Site; PPI=Protein-Protein Interaction.

### Analysis of differential gene – transcription factor networks

We conducted within-species comparisons of gene-TF networks which tested for differences between temperatures in *Chrysemys* (Cpi-FPT vs Cpi-MPT), while for *Apalone*, comparisons tested for differences between-sex per-temperature (Asp-F26 vs Asp-M26, Asp-F31 vs Asp-M31), and between-temperatures per-sex (Asp-F26 vs Asp-F31, Asp-M26 vs Asp-M31).

Networks consist of nodes connected by edges. An edge is a vertex connecting a gene to a TF, representing the presence of an interaction. We tested for edge weights that were significantly different between two networks of interest. Using a t-test, the difference between each predicted edge (e.g. network *i* edge minus network *ii* edge) was compared to the average difference for that edge between all permuted pairs of networks. Significance was assessed following a Benjamini-Hochberg correction for multiple comparisons.

In addition to examining differential network edges, we also examined differences in network targeting patterns, of which there are two types (Glass, et al. 2014; Glass, et al. 2015): gene in-degree and TF out-degree. Gene in-degree describes how many edges in the network point to a gene, while TF out-degree, describes how many edges point from a TF to various genes. Thus, these values help identify which genes and TFs of the networks are highly connected, and whether differences in the in-degree and out-degree patterns exist. In-degree and out-degree values were calculated with the pandaR function calcDegree() which sums edge weights for a particular vector (i.e., the row of TFs targeting one gene, or the columns of genes targeted by one TF). Similarly, for the differential edge calculation, a t-test was performed to assess differences in targeting patterns between pairs of empirical networks relative to the test distribution of differences in targeting patterns for all pairs of networks. Significance was assessed following a Benjamini-Hochberg correction for multiple comparisons.

We also assessed overall differences in the gene:TF networks by calculating the sum of squares of each network’s matrix and comparing the difference of the sum of squares between empirical networks to the distribution generated by calculating the differences in the sums of squares of all pairs of permuted networks. Thus, we assessed networks for differences at the edge (individual gene-by-TF targeting patterns), vector (TF regulatory patterns for collective gene targets), and matrix (whole network regulatory patterns) levels.

### Principal components analysis and trajectory analysis

We filtered all six networks (2 from *Chrysemys* and 4 from *Apalone*) to identify the set of genes and TFs shared between species and transformed negative edge-weights to zero to denote a lack of relationship. Principal components analysis was then performed on the edge weights representing the genes targeted by each TF. These networks were further filtered to only include TFs with an average expression level of >1.5 log_2_(TPM) across all libraries in the initial input gene expression data (to rule out negligibly expressed TFs) and that were present in all 6 networks. Next, the principal components were subjected to a modified trajectory analysis (Adams and Collyer 2009) to test for differences in the regulatory targeting patterns of orthologous TFs. This analysis connects centroids of orthologous TFs by a vector in regulome space. The centroid is the position of a TF in the regulome space informed by the genes it regulates in the networks within each species (and is calculated from each set of species-specific networks), and the distance between centroids measures how similar or different TFs are relative to one another in the genes they are predicted to regulate between species. Longer vectors denote greater divergence in the gene targets of ortholog TFs between species, compared to TFs connected by shorter vectors in the PC Euclidian plane. To measure these distances between species for each TF, we modeled gene targets as a function of TF, species, and their interaction (linear model: Gene Targets ∼ TF + Species + TF:Species). Distances between the centroids of the principal components of the target genes for a particular TF per species were calculated. We then generated a distribution of distances under the null hypothesis that there were no differences between species (no effect of species on TF). This corresponds to the prediction that no evolution has occurred in the regulatory role of these orthologous TFs since *Chrysemys* and *Apalone* split ∼175 mya (timetree.org) (hypothesis H0). We tested the distances of each TF between species against this distribution using a permutation test and applied a Benjamini-Hochberg correction for multiple comparisons with an FDR of 0.1. Using this level of FDR the top ∼5% of distances of TFs and those belonging to the right-most distribution in our bimodal distribution of distances were considered significant. Non-significant distances between orthologous TFs between species reject hypothesis H1 and may denote TF hubs conserved between lineages in their target patterns (supporting hypothesis H0), whereas orthologous TF hubs with significantly longer distances would have diverged between species in the identity of the genes they regulate (hypothesis H1) (Fig 1). We tested hypothesis H2 by measuring trajectory distances of non-orthologous pairs of TFs between species. Here we identified interspecific trajectories that were shorter than expected under the null hypothesis, indicating non-orthologous (analogous) TFs that converged on the same gene targeting pattern between species (supporting hypothesis H2). Additionally, we confirmed that these trajectories were shorter than the distance to their respective orthologous TFs (albeit not significantly shorter since they were all above the critical value of the 5% quantile). Hypothesis H3 is tested by consilience with the other hypotheses.

### Gene ontology overrepresentation analysis of networks

The PANTHER (v17.0 – date: 2022-02-02) online GUI (Thomas, et al. 2022) was used to conduct overrepresentation analysis of gene targets of TFs of interest. We used a Fisher’s Exact Test with an FDR correction to determine which gene ontology (GO SLIM) terms (molecular function, cellular component, and biological process) were overrepresented in the targets(Mi, et al. 2019), as well as if any PANTHER pathways (Mi and Thomas 2009) or PANTHER protein classes were overrepresented. Since our focal species are non-model organisms that are not represented in the PANTHER databases, we first mapped the translated CDSs from *Chrysemys* to PANTHER IDs using PANTHER HMM scoring tools (pantherScore2.2) to score our sequences against the PANTHER HMM library (v17.0), following the instructions provided by the PANTHER developers. In the case of duplicate hits due to redundancy in the *Chrysemys* genome, we prioritized the result to those with the highest bitscore. Since *Apalone* annotations were based on the *Chrysemys* genome, we used the mappings obtained from *Chrysemys* CDS sequences to transfer the corresponding annotations to genes in the *Apalone* networks. Only TFs with at least 50 known gene targets prior to uploading to PANTHER were included in the overrepresentation analysis, as functional annotation results can be sensitive to input list size (Zhao and Rhee 2023; Ziemann, et al. 2024). Additionally, to focus the analysis on gene targets with the highest support in the network models, gene targets were filtered to retain those with an edge weight in the top 5% of all edge weights. Our approach resulted in sets exceeding the 50-gene benchmark, as most sets analyzed had 100s or 1000s gene targets in their input list. Tabular results of the analysis were downloaded and saved locally. The background gene set included all genes present in the network, but because not all genes mapped to panther IDs the total effective number was smaller (CPI: 16489 genes, ASP: 11386 genes).

### Transcription factor functional similarity analysis via calculation of semantic similarity

The resulting sets of overrepresented terms of TF gene targets were compared for semantic similarity with GOGO (Zhao and Wang 2018) to determine their degree of functional similarity. GOGO is a hybrid algorithm for semantic similarity calculations that uses both the topology of the directed acyclic graphs (DAG) that make up the ontology (which informs ancestor-child term relationships) and considers the number of children nodes of a term (which reveals the information contained in a term). Thus, it allows us to impartially assess how similar two lists of gene ontology terms are to one another which can be hindered by their hierarchical nature. We updated the DAGs used by the program to match the same reference ontologies used to run the overrepresentation tests (PANTHER v17.0). We used the gene_list_comb.pl script to calculate semantic similarity of the sets of statistically significant GO terms returned for the gene targets of a TF in a particular network. We then compared the similarity scores, which range from zero (no similarity) to one (perfect overlap), to assess the degree of functional similarity of TF targets for orthologous TFs across networks. We used a Mann-Whitney U test to evaluate whether there were significant differences in semantic similarity for the set of within-species comparisons (all *Chrysemys* by *Chrysemys* and all *Apalone* by *Apalone* contrasts) relative to the set of between-species comparisons (all *Chrysemys* by *Apalone* contrasts) and applied a Bonferroni correction for multiple comparisons.

### Subnetworks

Because the final networks obtained from PANDA for *Chrysemys* contained 2,883,632 edges, we also generated subnetworks to focus our attention on the most strongly supported edges, by filtering networks for the edges with an arbitrary selected Z score > 10 (which correspond to the top 0.03% of edges for *Chrysemys* and *Apalone*). Subnetworks were visualized in Cytoscape (v3.9.0)(Shannon, et al. 2003), and the hubs and their targets were identified using the function targetedGenes() for overrepresentation analysis of gene ontology functional terms.

All scripts used for the analyses are included in the Supplementary Materials.

## Acknowledgements

We thank Parnal Joshi for help in preparing the protein-protein interaction dataset used in the generation of the networks. We thank Leila Fattel for analytical suggestions in our functional annotation. We note that ChatGPT was used to help brainstorm some R commands in order to streamline code generation, and any resulting code was then tested and fine-tuned by TBG for validation to ensure it achieved the appropriate aim.

This work was funded in part by US NSF grants IOS 1555999 and IOS 2127995 to NV. The research reported in this paper is partially supported by the HPC@ISU equipment at Iowa State University, some of which has been purchased through funding provided by NSF under MRI grants number 1726447 and 2018594. All opinions, findings, and conclusions expressed in this paper are those of the authors.

## Data Availability Statement

No new data were generated for this paper and have been previously published; however sequencing reads are available in the Short Read Archive at NCBI: BioProjects PRJNA683586 (*Apalone spinifera* embryos—SRR13224849-SRR13224888) and PRJNA594037 (*Chrysemys picta* embryos—study number SRP237291; SRR10674595-SRR10674614).

## Abbreviations

CaRe: calcium redox
DAG: directed acyclic graph
ESD: environmental sex determination
FPT: female producing temperature
GRN: gene regulatory network
GSD: genotypic sex determination
MPT: male producing temperature
Mya: million years ago
PC: principal components
ROS: reactive oxygen species
SDM: sex determination mechanism
TF: transcription factor
TFBS: transcription factor binding site
TSD: temperature-dependent sex determination

## Notes

### Competing Interest Statement

The authors have declared no competing interest.

